# A supracellular actin network transmits forces over long distances at the apical surface of squamous carcinoma cells

**DOI:** 10.1101/2025.01.10.632450

**Authors:** Léa Marpeaux, Claire Baudouin, Lara Elis Alberici Delsin, Cédric Plutoni, Gregory Emery

**Author notes:** These authors contributed equally to this work (Léa Marpeaux, Claire Baudouin).

## Abstract

Epithelial tissues form protective barriers while supporting critical functions such as absorption and secretion. Their structural and functional integrity relies on adherens junctions, which coordinate migration and transmit forces between adjacent cells by connecting their actin cytoskeleton. In this study, we report the presence of an apical supracellular actin network in squamous epithelial cells. Using squamous carcinoma A431 cells as a model, we characterized this network composed of star-shaped actin structures interconnected by linear actin bundles that span multiple cells. We demonstrate that the network’s formation and maintenance require actomyosin contractility and intact adherens junctions, while tight junctions seem dispensable. Furthermore, this network dynamically reorganizes as cells migrate and preferentially aligns with the direction of movement. This contractile structure generates mechanical tension that extends across the apical surface of multiple cells. Our findings suggest that this supracellular actin network functions as a long-range force transmission device in squamous cells, advancing our understanding of the biomechanical properties of epithelia.

## INTRODUCTION

Multicellular organisms rely on the intricate organization of diverse cell types into specialized tissues, a process fundamental to embryonic development and tissue homeostasis. Among the various tissue types, epithelial tissues are of paramount importance. By covering both the internal and external surfaces of organisms, epithelia form a protective barrier against environmental factors while also playing essential roles in absorption and secretion (Marchiando et al., 2010).

Epithelial tissues consist of tightly connected cells and are classified based on cell shape and the number of layers they form. For instance, squamous epithelia are composed of large and flat cells arranged into single or multiple layers to form simple or stratified epithelia, respectively (Lemke and Nelson, 2021). To ensure their functions, epithelial cells require an apical-basal polarity. This polarity is maintained by various cell-cell junctions, mainly tight junctions at the apico-basolateral border and adherens junctions along the basolateral membrane (Buckley and St Johnston, 2022). These intercellular junctions are essential for maintaining tissue integrity, regulating transepithelial transport and promoting coordinated cellular responses (Tunggal et al., 2005, Markov et al., 2010, Arboleda-Estudillo et al., 2010).

Adherens junctions, composed of cadherin transmembrane proteins and associated intracellular catenins (Ozawa et al., 1989, Reynolds et al., 1994), play a crucial role in linking the actin cytoskeletons of adjacent cells (Rimm et al., 1995, Ozawa et al., 1989, Reynolds et al., 1994). When non-muscle myosin II motor proteins couple with actin filaments, contractile forces are generated (Peterson et al., 2004) leading to the recruitment of mechanosensitive proteins like vinculin to the adherens junctions (Yonemura et al., 2010). This recruitment strengthens the junctions (Thomas et al., 2013) and facilitates force transmission and mechanical communication between neighboring epithelial cells (Borghi et al., 2012, Cai et al., 2014). At the tissue level, these intercellular adhesions serve as relays to build supracellular actomyosin structures that extend over multiple cells (Williams-Masson et al., 1997, Danjo and Gipson, 1998, Cheng et al., 2004, Laplante and Nilson, 2006, Martin et al., 2009, Monier et al., 2010, Röper, 2012, Wang et al., 2020).

Supracellular actomyosin structures have been described in various processes. For example, several studies have highlighted the importance of a supracellular actomyosin ring at the leading edge of wounded epithelia (Bement et al., 1993, Danjo and Gipson, 1998, Wood et al., 2002, Tamada et al., 2007). This basal structure, observed in many organisms, contributes to wound closure through contractility, acting as a purse-string (Martin and Lewis, 1992, Bement et al., 1993, Danjo and Gipson, 1998). Other studies have demonstrated the importance of supracellular actin structures in maintaining a hierarchical organization within migrating epithelial cells. This hierarchy coordinates actively protruding leader cells, that guide the movement, with follower cells to form a cohesive unit. For instance, we and others described a supracellular actomyosin cortex surrounding Drosophila border cell clusters which restricts protrusion formation to the leader cell and promotes directional migration (Ramel et al., 2013, Wang et al., 2020). Basal supracellular actomyosin arcs identified in cultured Madin-Darby canine kidney (MDCK) epithelial cells were shown to inhibit protrusions in some cells at the migration front, ensuring that only a limited number of cells lead the migration (Fenteany et al., 2000, Reffay et al., 2014). Moreover, apical supracellular actin structures have been observed in mouse skin keratinocytes (Vaezi et al., 2002). However, their nature and function remain elusive as no subsequent characterization has been performed, possibly due to the lack of an easily accessible model.

In this study, we report the discovery of an apical supracellular actin network (ASAN) that extends across multiple normal and cancerous squamous cells migrating collectively. ASAN shares similarity with the supracellular actin observed in skin keratinocytes but is more extended. We characterized this network in migrating squamous carcinoma A431 cells. It is composed of linear bundles of actin filaments spanning the entire apical cell surface and star-like actin structures connecting the bundles present in surrounding cells. Further observations revealed that ASAN preferentially aligns with the direction of migration and undergoes continuous reorganization throughout the migration process. Interestingly, the formation and maintenance of this network depend on contractility and intact adherens junctions, while tight junctions seem dispensable for its assembly. Importantly, we discovered that this network, which is contractile and under tension, exerts forces on the apical cell surface at a multicellular level. Overall, our data indicate that ASAN functions as a long-range force transmission device in skin epithelial cells providing new insights into the mechanical properties of epithelial tissues.

## RESULTS

### Collective migration of normal and cancerous epithelial cells results in finger formation with mesenchymal-like cells at their tips

To investigate the collective migration patterns of both normal and cancerous human cell lines from various tissues, we used a standardized 2D wound-healing assay. Cells were cultured on glass substrates and confined in a silicon insert with two wells separated by a precise 500 µm gap. Once the cells formed confluent sheets, the insert was carefully removed, triggering spontaneous cell migration into the surrounding environment.

We acquired bright-field images of the periphery of migrating cell sheets at 24 h intervals for each cell line (Fig. 1A and B and Fig. 2 A-J). This approach enabled quantification of diverse migration parameters over a 72 h period following insert removal.

**Figure 1.**
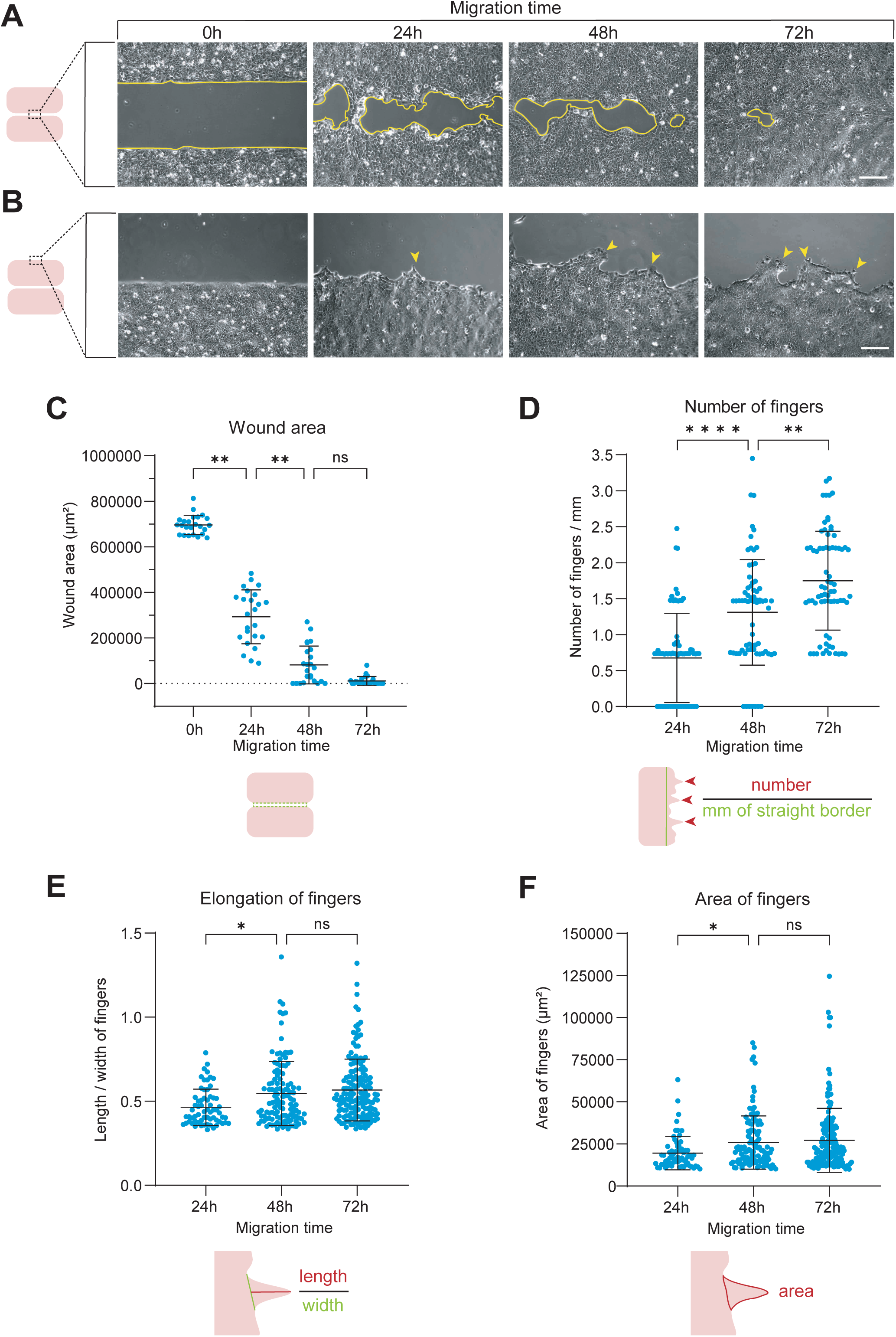
A431 skin squamous carcinoma cells form fingers with faster-moving mesenchymal-like cells at their tips during sheet migration. **(A)** Representative bright-field images of the gaps between both A431 cell sheets at 0, 24, 48 and 72h of migration after removing the insert. The yellow lines represent the contour of the gaps not covered by the cells (wound area). Scale bar = 200 µm. **(B)** Representative bright-field images of the outside periphery of both A431 cell sheets at 0, 24, 48 and 72h of migration after removing the insert. The yellow arrowheads point towards multicellular structures called fingers. Scale bar = 200 µm. **(C)** Wound area after 0, 24, 48 and 72h of A431 cell migration. For each condition, 8 pictures per experiment obtained in 3 independent experiments were analysed. **(D-F)** Mean number of fingers/mm **(D)**, elongation of fingers **(E)** and area of fingers **(F)** formed after 24, 48 and 72h of A431 cell migration. The elongation was calculated by the ratio length/width of fingers and the area by the entire surface over the base of the fingers. For each condition, 24 pictures per experiment obtained in 3 independent experiments were analysed. All the data are presented as mean ± standard deviation (s.d.) and tested by Kruskal-Wallis (ns: not significant; *: p<0,05; **: p<0,01; ****: p<0,0001).

**Figure 2.**
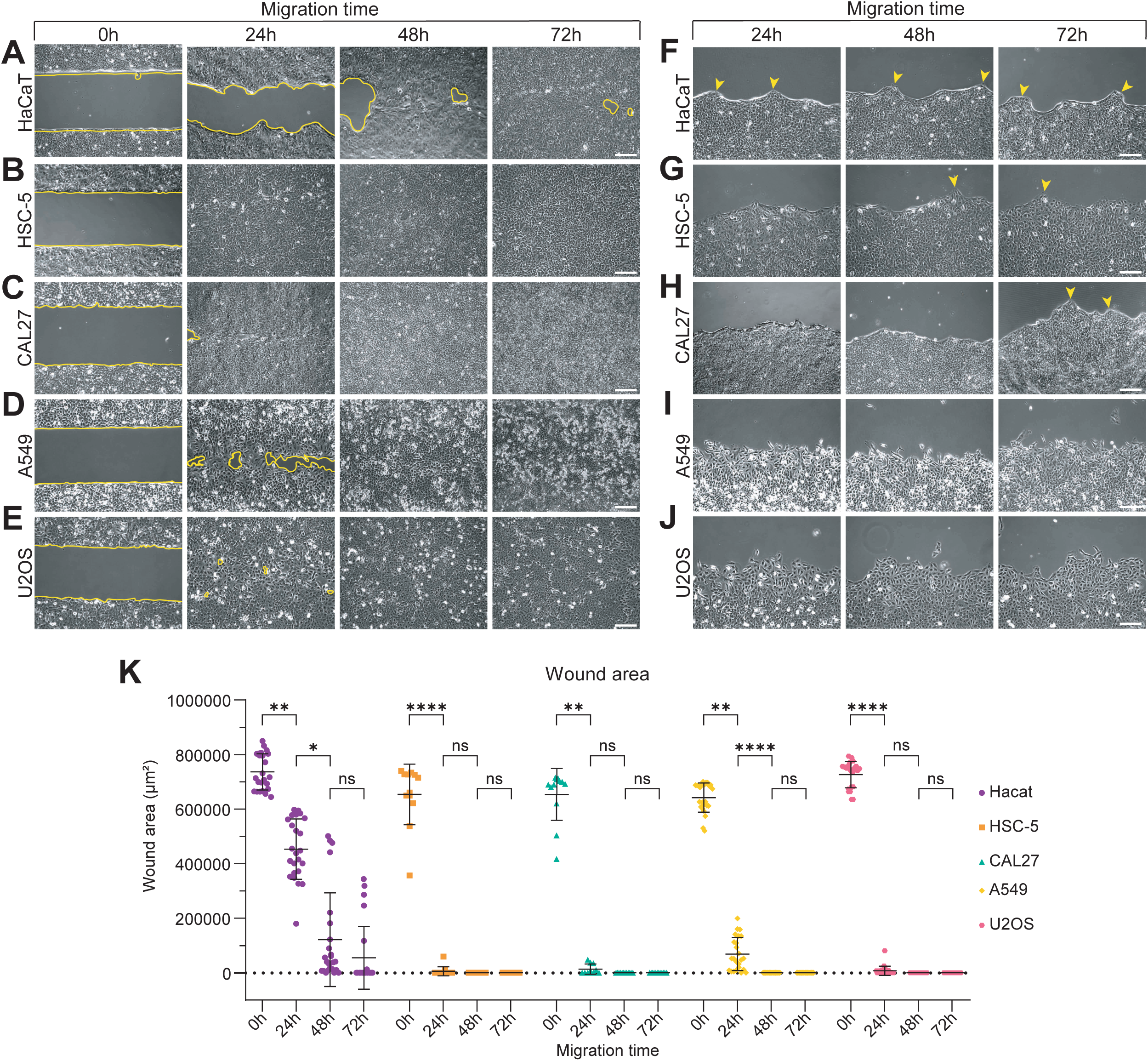
HaCaT, HSC-5 and CAL27 epithelial cells form fingers when migrating collectively but not A549 and U2OS cells. **(A-E)** Representative bright-field images of the gaps between both cell sheets of HaCaT **(A)**, HSC-5 **(B)**, CAL27 **(C)**, A549 **(D)** and U2OS **(E)** cell sheets at 0, 24, 48 and 72h of migration after removing the insert. The yellow lines represent the contour of the gaps not covered by the cells (wound area). Scale bar = 200 µm. **(F-J)** Representative bright-field images of the outside periphery of both HaCaT **(F)**, HSC-5 **(G)**, CAL27 **(H)**, A549 **(I)** and U2OS **(J)** cell sheets at 0, 24, 48 and 72h of migration after removing the insert. The yellow arrowheads point towards multicellular structures called fingers. Scale bar = 200 µm. **(K)** Wound area after 0, 24, 48 and 72h of HaCaT, HSC-5, CAL27, A549 and U2OS cell migration. For each condition, 3 to 8 pictures per experiment obtained in 3 independent experiments were analysed. All the data are presented as mean ± s.d. and tested by Kruskal-Wallis (ns: not significant; *: p<0,05; **: p<0,01; ****: p<0,0001).

The closure of the gap between both cell sheets was monitored by measuring the wound area. A431 skin squamous carcinoma cells covered the defined area within 48 to 72 h (Fig. 1A and C). The outer periphery of the cell sheets revealed the formation of hierarchical structures, called “fingers” with one or more protruding leader cells at the tip, followed by a stream of follower cells (Fig. 1B). Since A431 cells formed prominent fingers, we characterized these multicellular structures by quantifying their number, their elongation through their length-to-width ratio and their area. A431 cells showed a significant increase in the number of fingers between 24 and 72 h of migration (Fig. 1B and D). Regarding the elongation and area of fingers, they also increased significantly between 24 and 48 h (Fig. 1B, E and F). Like A431 cells, HaCaT epidermal keratinocytes closed the wound within 48 to 72h (Fig 2A and K). Both cell lines migrated slower than HSC-5 skin squamous carcinoma cells, CAL27 tongue squamous carcinoma cells, A549 lung adenocarcinoma cells and U2OS osteosarcoma cells which nearly or entirely closed the gap within 24 h (Fig. 2B-E and K).

HaCaT cells, and to a lesser extent HSC-5 and CAL27 cells, formed finger-like structures, in a similar manner as A431 cells (Fig. 2F-H). In contrast, A549 and U2OS cells migrated collectively without forming such structures (Fig. 2I and J). These results suggest that the hierarchical “finger-like” migration, previously observed in MDCK cells (Poujade et al., 2007, Petitjean et al., 2010, Reffay et al., 2011, Reffay et al., 2014, Yamaguchi et al., 2015, Balcioglu et al., 2020, Isozaki et al., 2020), might be a common thread of normal and cancerous epithelial cells. However, this migration pattern does not appear to predict the migration speed of cell collectives (Fig. 1B and C and Fig. 2F-K).

As A431 cells form the most prominent finger-like structures compared to the other cell lines, we decided to investigate their finger shape changes over time. To do so, we compared the dynamics of protrusion and retraction of leader cells at the tip of the fingers to those of cells on their sides. A kymograph generated over a period from 56 to 60 h of migration revealed that leader cells undergo repeated cycles of protrusion extension and retraction, while side cells remain relatively steady (Fig. 3A). Moreover, velocity measurements indicated that leader cells began to move faster than side cells after 32 h of migration, with the velocity of leader cells increasing over time while the velocity of side cells remained stable (Fig. 3B). This significant difference in velocity between both cell types explains the finger growth and extension observed between 24 and 72 h of migration (Fig. 1B and D-F).

**Figure 3.**
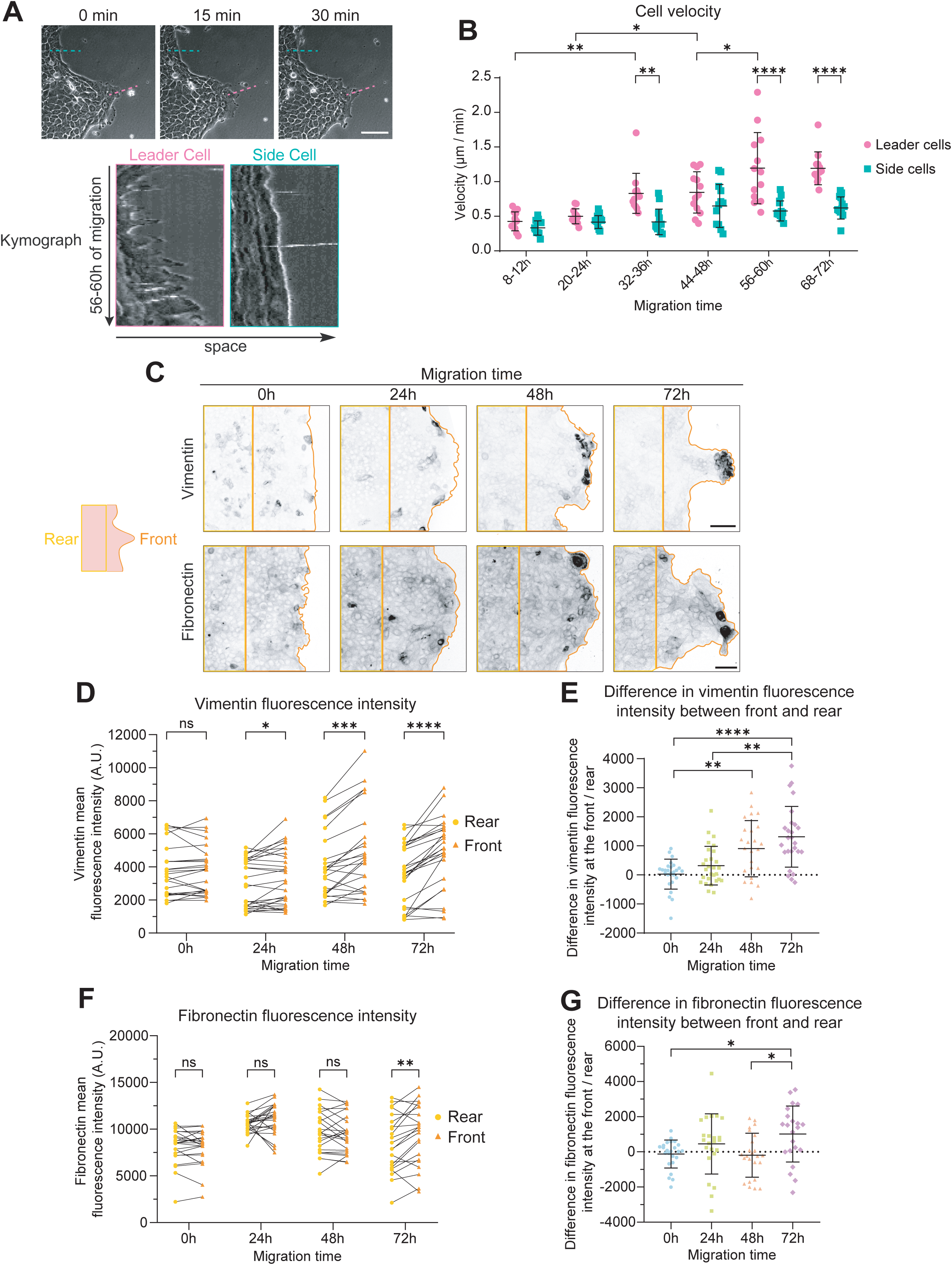
A431 skin squamous carcinoma cells form faster-moving mesenchymal-like cells at their tips during sheet migration. **(A)** Time-lapse DIC imaging of a finger formed by A431 cells after 56h of migration (top). Three images took at 15 min intervals show the protrusion and retraction dynamics of cells at the front of migration. Scale bar = 100 µm. Two kymographs, generated by drawing lines in the middle of a leader cell (pink line, left rectangle) and a side cell (blue line, right rectangle), show the protrusion and retraction dynamics of both cell types from 56 to 60h of migration (bottom). **(B)** Mean velocity of leader cells and side cells at different timepoint intervals during A431 cell migration. For each time interval, 12 leader cells and 12 side cells from one experiment were analysed. **(C)** Z-projections of representative confocal images of A431 cell sheets stained for vimentin and fibronectin after 0, 24, 48 and 72h of migration. Orange and yellow lines delimit respectively the front and rear of cell sheets. Scale bar = 100 µm **(D)** Vimentin mean fluorescence intensity at the front and at the rear of A431 cell sheets after 0, 24, 48 and 72h of migration. For each condition, 7 to 10 pictures per experiment of 3 independent experiments were analysed. **(E)** Difference in vimentin mean fluorescence intensity between the front and the rear in A431 cell sheets after 0, 24, 48 and 72h of migration. 7 to 10 pictures per experiment of 3 independent experiments were analysed. **(F)** Fibronectin mean fluorescence intensity at the front and at the rear of A431 cell sheets after 0, 24, 48 and 72h of migration. For each condition, 7 to 8 pictures per experiment of 3 independent experiments were analysed. **(G)** Difference in fibronectin mean fluorescence intensity between the front and the rear in A431 cell sheets after 0, 24, 48 and 72h of migration. 7 to 8 pictures per experiment of 3 independent experiments were analysed. All the data are presented as mean ± standard deviation (s.d.; B, E, G) or individual data points (D, F) and tested by Two-way ANOVA (B), Wilcoxon (D, F) or Kruskall-Wallis (E, G) (ns: not significant; *: p<0,05; **: p<0,01; ***: p<0,001; ****: p<0,0001).

To further elucidate the increased motility of leader cells, we examined the distribution of vimentin and fibronectin, two epithelial-to-mesenchymal transition (EMT) markers associated with enhanced migratory properties of leading-edge cells in vitro and in vivo (Dulbecco et al., 1983, Boyer et al., 1989, Walker et al., 2007, Walker et al., 2010, Menko et al., 2014, Bleaken et al., 2016, LeBert et al., 2018, Summerbell et al., 2020). Confocal microscopy revealed that vimentin and fibronectin were initially randomly distributed throughout the A431 cell sheet (Fig. 3C). As migration progressed, vimentin and fibronectin expression became restricted to the migration front, mainly in leader cells (Fig. 3C). This restriction led to a significant difference in vimentin intensity between the front and rear of the cell sheet starting from 24 h of migration and increasing over time (Fig. 3D-E). Quantification of fibronectin intensity also revealed a significant difference in expression between the front and rear of the cell sheet at 72 h of migration (3F-G). These observations suggest that leader cells at the tip of the migratory fingers might undergo a partial EMT, allowing them to migrate faster than cells at the back while remaining physically connected to them, and contributing to the formation and growth of fingers during migration.

### Collectively migrating skin epithelial cells show a dense and apical supracellular actin network

Finger formation during collective cell migration supposedly relies on a tight communication between leader cells and their followers. Supracellular actin structures have been described to facilitate intercellular communication across various collective cell migration models (Danjo and Gipson, 1998, Jacinto et al., 2002, Köppen et al., 2006, Hidalgo-Carcedo et al., 2011, Reffay et al., 2014, Plutoni et al., 2019). To elucidate the role of actin in the finger-like migration, we examined filamentous actin at the periphery of the cell sheets for the different cell lines after 72 h of migration using confocal microscopy. Distinct Z-projections for both basal and apical planes were generated (Fig. 4A and B and Supplementary Fig. 1A-E).

**Figure 4.**
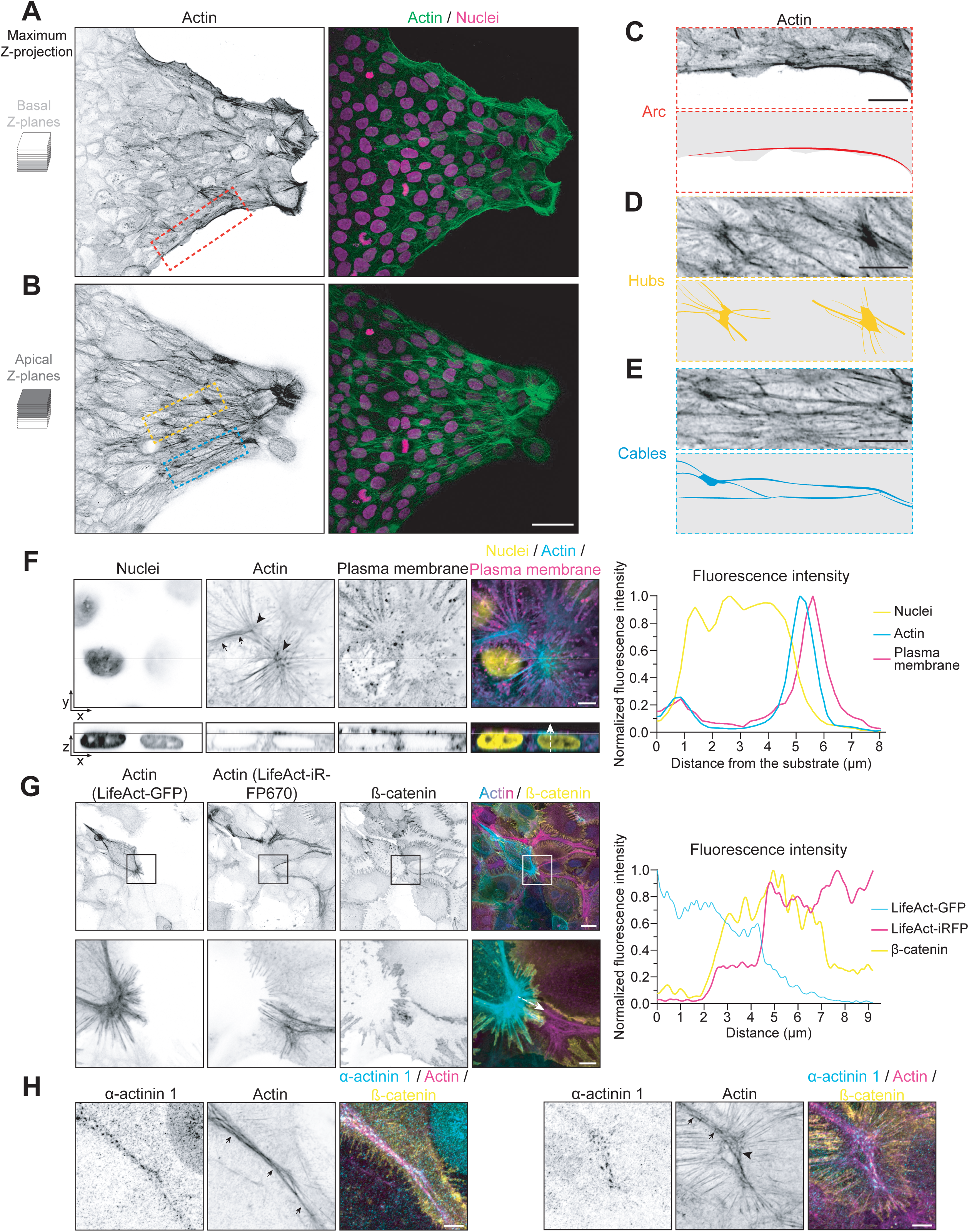
Collectively migrating A431 cells contain an apical supracellular actin network (ASAN) **(A-B)** Representative maximum intensity projections of the basal (A) and apical (B) Z-planes of a finger formed after 72h of A431 migration. The actin and nuclei were stained. The red, yellow and blue rectangles show respectively the actin arcs, hubs and cables. Scale bar = 50 µm. **(C-E)** Magnifications of the red, yellow and blue rectangles from the previous images showing areas with different actin structures: arcs (C), hubs (D) and cables (E). Each panel displays the actin signal (top) and a schematic representation (bottom) of the structure. Scale bar = 20 µm. **(F)** Representative xy (top) and xz (bottom) images at enhanced resolution acquired with an Airyscan detector, of a confocal Z-stack (left) showing an area with hubs (black arrowheads) and cables (black arrows) in A431 cells after 72h of migration. A white arrow was drawn from the substrate to the top on the xz merge image and used to generate a plot profile (right) representing the normalized fluorescence intensity of each signal along the arrow. Scale bar = 5 µm. **(G)** Representative maximum intensity projections of the apical Z-planes showing an area with cables in A431 cells after 72h of migration (left). A mosaic of cells stably expressing the filamentous actin probe LifeAct fused to either GFP or iRFP670 was generated and stained for β-catenin. Full images (top) and magnifications (bottom) are shown. Magnifications are high resolution images captured using an Airyscan detector. A white arrow was drawn from a LifeAct-GFP expressing cell to a LifeAct-iRFP670 expressing cell and used to generate a plot profile (right) representing the normalized fluorescence intensity of each signal along the arrow. Scale bar = 20 µm (top) and 5 µm (bottom). **(H)** Representative maximum intensity projections of the apical Z-planes showing a cable (arrows, left) and a hub linked to a cable (arrowhead and arrows, right) in A431 cells after 72h of migration. α-actinin 1, actin and β-catenin were stained. These are high resolution images captured using an Airyscan detector. Scale bar = 5 µm

Z-projections of the basal planes revealed predominantly cortical actin and stress fibers in all the tested cell lines (Fig. 4A and Supplementary Fig. 1A-E). However, A431, HaCaT, HSC-5 and CAL27 cells also displayed actin arcs along the sides of the fingers (Fig. 4A and C and Supplementary Fig. 1A-C). These arcs, previously undescribed in these cells to our knowledge, are morphologically identical to the basal supracellular actin structures that inhibit protrusion formation in various in vitro and in vivo collective cell migration models (Bement et al., 1993, Kiehart et al., 2000, Panfilio and Roth, 2010, Reffay et al., 2014, Czerniak et al., 2016).

Surprisingly, Z-projections of the apical planes showed that A431, HaCaT and HSC-5 cells, which are normal and cancerous epithelial cells of skin origin, exhibited a complex actin network (Fig. 4B and Supplementary Fig. 1A and B). In these cell lines, the actin network was composed of two main structures connected to each other: hubs and cables. Hubs appeared as actin stars with many actin filaments or bundles radiating from a central spot, while cables were long actin bundles of varying thickness (Fig. 4B, D and E and Supplementary Fig. 1A and B). The density of the network varied among these three cell lines, with A431 and HSC-5 cells exhibiting a denser network compared to HaCaT cells (Fig. 4B and Supplementary Fig. 1A and B). In contrast, CAL27 non-skin epithelial cells showed hubs and actin filaments but prominent cables were found to be rare (Supplementary Fig. 1C). Non-epithelial A549 and U2OS cells showed only cortical actin without the formation of such network (Supplementary Fig. 1D and E). These findings suggest that normal and cancerous squamous epithelial cells are more prone to form an apical actin network. However, the role of this network in the hierarchical migration of cells is still undetermined.

Given that such an apical actin organization has not been previously described to our knowledge, we further characterized this network. To do so, we focused our study on the dense actin network observed in A431 cancer cells. Nuclear and plasma membrane stainings confirmed the apical localization of the actin network (Fig. 4F). A plot profile showed that it was restricted above the nuclei, close to the apical plasma membrane (Fig. 4F).

To determine individual cell contributions to the actin network, we generated a mosaic of migrating A431 cells stably expressing the filamentous actin probe LifeAct fused either to GFP or iRFP670. Staining of this mosaic for β-catenin allowed us to visualize adherens junctions (Piepenhagen and Nelson, 1993). We observed that multiple cells contributed to the actin network by forming cables or hubs that interconnected, creating a continuous structure (Fig. 4G). A plot profile demonstrated the transition between a GFP-labeled cable from one cell to an iRFP670-labeled cable from another cell (Fig. 4G). Moreover, the connection between both cables was marked by the presence of β-catenin (Fig. 4G). This indicates that the actin network is supracellular and suggests that it is connected through adherens junctions. Given the apical nature of the supracellular actin network in A431 cells, we decided to call it ASAN for “Apical Supracellular Actin Network”.

The high density of some actin hubs and cables led us to hypothesize that ASAN might be enriched with actin-bundling proteins, particularly α-actinin-1 (Maruyama and Ebashi, 1965). This ubiquitously expressed protein is mainly known for its role in bundling parallel actin filament to form actin stress fibers (Lazarides and Burridge, 1975, Kovac et al., 2013, Feng et al., 2013). Immunofluorescence experiments in migrating A431 cells revealed that α-actinin-1 formed well-defined punctae within specific cables and hubs (Fig. 4H). This distribution pattern suggests a dual role for α-actinin-1 in the actin network: it may bundle actin filaments parallel to each other along cables while potentially linking actin cables in various orientations within hubs. However, since α-actinin-1 was not present in every structure of ASAN, it is likely that other actin-bundling proteins might also participate in the formation of the network.

### ASAN is remodelled during A431 collective cell migration

While we previously characterized ASAN in A431 cells after 72 h of migration, we next sought to understand its formation and reorganization throughout the collective cell movement.

Analysis of the apical actin organization over time revealed that a network with cables and hubs was already present at the onset of migration (Fig. 5A). However, this network underwent significant changes throughout the migration process (Fig. 5A). Quantitative analysis showed important changes in the distribution and characteristics of these structures over time (Fig. 5B-D). Specifically, the number of hubs significantly decreased during the first 24 h of migration, followed by a recovery to initial levels (Fig. 5B). In contrast, both the area of hubs and the total length of cables exhibited significant increases after 48 and 24 h, respectively (Fig. 5C and D). These last changes contribute to the apparent densification of ASAN during cell migration, which presents significantly more supracellular actin structures, particularly after 24 h (Fig. 5A-D).

**Figure 5.**
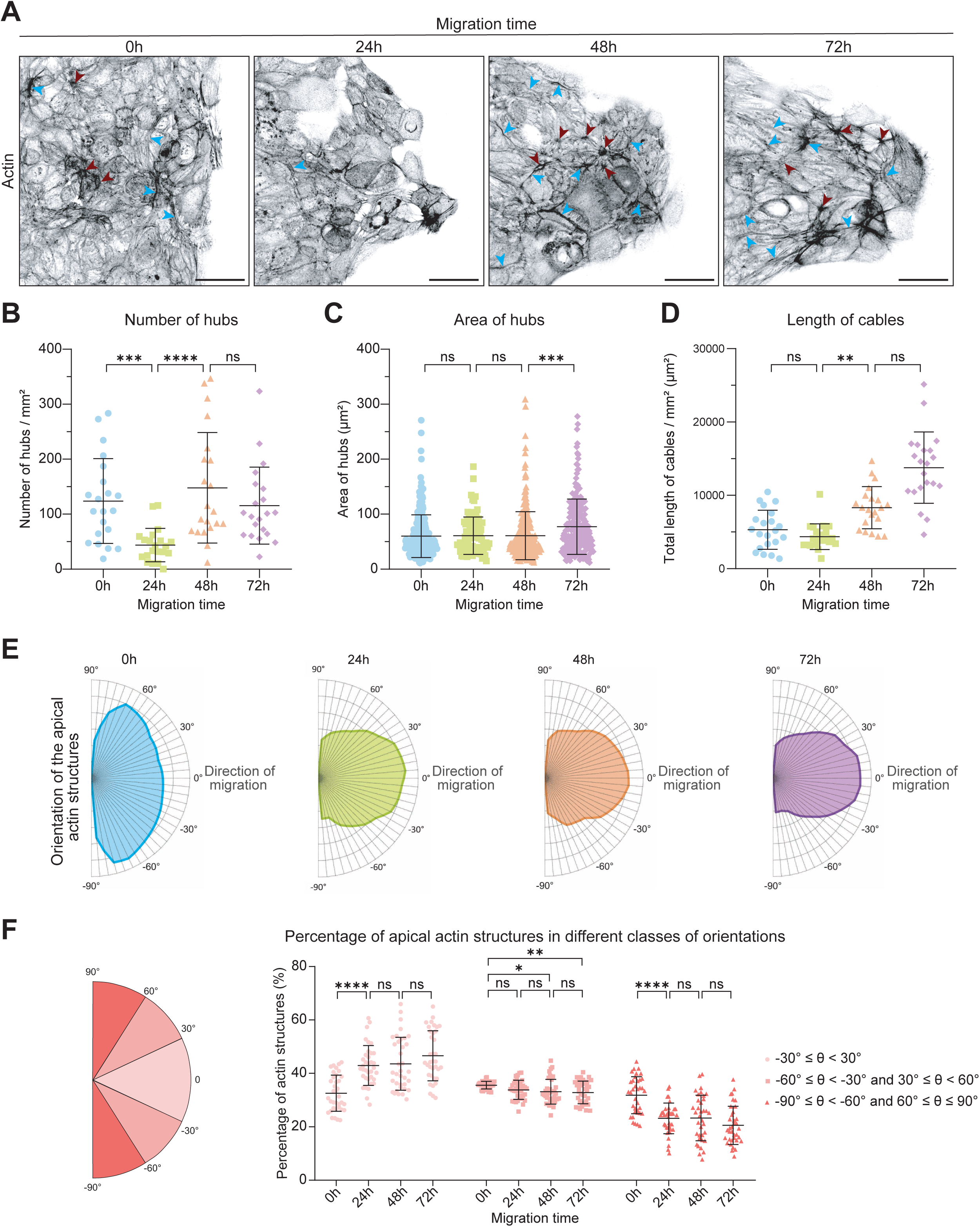
The apical supracellular actin network remodels and reorientates during A431 collective migration. **(A)** Representative maximum intensity projections of the apical Z-planes of the A431 cell sheet periphery after 0, 24, 48 and 72h of migration and actin staining. Scale bar = 50 µm. Red and blue arrowheads point towards hubs and cables. **(B-D)** Number of hubs per mm² (B), area of hubs (C) and total length of cables per mm² (D) after 0, 24, 48 and 72h of A431 cell migration. 7 pictures per experiment of 3 independent experiments were analysed. **(E)** Mean orientation of the apical actin structures after 0, 24, 48 and 72h of A431 cell migration. The orientation goes from -90° to 90° with 0° corresponding to the direction of migration. 12 pictures per experiment of 3 independent experiments were analysed. **(F)** Percentage of apical actin structures after 0, 24, 48 and 72h of A431 cell migration with angles from -30° to 30°, angles from -60° to -30° and from 30° to 60° and angles from -90° to -60° and from 60° to 90°. 12 pictures per experiment of 3 independent experiments were analysed. All the data are presented as mean ± s.d. and tested by Kruskal-Wallis (B-D) or Two-way ANOVA (F) (ns: not significant; **: p<0,01; ***: p<0,001; ****: p<0,0001)

In addition to changes in ASAN density, we observed a marked shift in its orientation during the migration process. Initially, the apical actin structures were randomly oriented whereas, starting from 24 h, they became preferentially aligned with the direction of migration, defined as an angle of 0° (Fig. 5E and F). Quantitative analysis showed a significant increase in the percentage of actin structures oriented between -30° and 30°, coupled with a significant decrease in the percentage of actin structures oriented between - 90° to -60° and 60° to 90° from 0 to 24 h of migration (Fig. 5F). Moreover, the percentage of actin structures oriented between -60° to -30° and 30° to 60° decreased throughout the migration (Fig. 5F). These results show that ASAN undergoes a major reorientation within the first 24 h of migration and continues to refine this alignment as migration progresses. This reorientation precedes densification. Further investigation will be needed to determine whether the reorientation of ASAN during cell migration is a cause or a consequence of the directionality of migration.

### Adherens junctions are essential for the formation of ASAN in A431 cells

Supracellular actin structures often depend on adherens junctions for their formation (Williams-Masson et al., 1997, Danjo and Gipson, 1998, Cheng et al., 2004, Laplante and Nilson, 2006, Martin et al., 2009, Monier et al., 2010, Röper, 2012, Wang et al., 2020). Moreover, our mosaic experiment revealed the presence of β-catenin at the junctions between actin cables and hubs from adjacent cells (Fig. 4G). This suggests that cadherin-based adherens junctions may connect individual actin structures to assemble ASAN.

To test this hypothesis, we treated A431 cells, stably expressing a LifeAct fluorescent actin probe, with 4 mM EGTA after 72 h of migration and conducted time-lapse imaging. EGTA is a calcium chelator commonly used to destabilize adherens junctions by disrupting calcium-dependent cadherin interactions (Volberg et al., 1986). As expected, we observed a decrease in fluorescent E-cadherin levels at the junctions after EGTA treatment compared to the control (Fig. 6A and Movies 1 and 2). Furthermore, within approximately 20 min, ASAN completely disappeared, while it remained intact in control cells (Fig. 6A and Movies 1 and 2). Notably, disruption of the apical actin network began as early as 4 min post-EGTA treatment, suggesting that adherens junctions might be critical for maintaining ASAN in A431 cells. Interestingly, this process was reversible as the washout of EGTA treatment induced the reappearance of ASAN within a few hours (Supplementary Fig. 2A and Movie 3). This indicates that adherens junctions might also be essential for the formation of ASAN in A431 cells.

**Figure 6.**
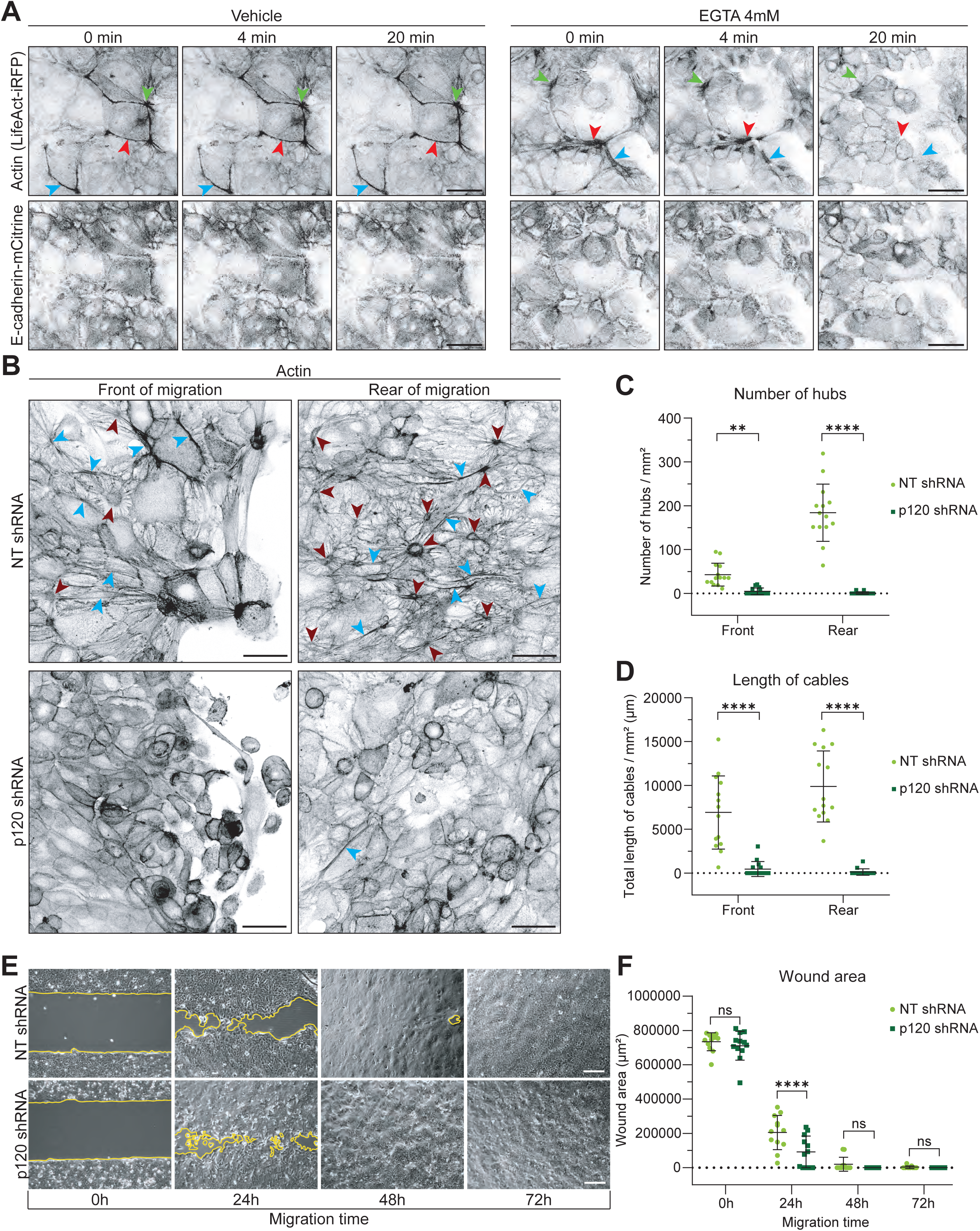
Adherens junctions are necessary for the formation and maintenance of ASAN and the efficient collective migration of A431 cells. **(A)** Representative maximum intensity projections of the apical Z-planes of a part of an A431 cell sheet that migrated during 72h before being treated with the vehicle (control, left) or 4 mM of the calcium chelator EGTA (right). 0 min corresponds to the beginning of the treatment. Cells are stably expressing the actin probe LifeAct-iRFP670 and the adherens junction protein E-cadherin-mCitrine. Green, red and blue arrowheads point towards cables and hubs that are maintained in the control condition but disappear after EGTA treatment. Scale bar = 50 µm. **(B)** Representative maximum intensity projections of the apical Z-planes of the front (left) and rear (right) of migration of A431 cell sheets. Using non-target and specific shRNAs, control (top) and p120-catenin (bottom) knocked down cells were generated, allowed to migrate during 72h and stained for actin. Red and blue arrowheads point towards hubs and cables. Scale bar = 50 µm. **(C, D)** Number of hubs per mm² (C) and total length of cables per mm² (D) at the front and at the rear of migration of control and p120-catenin knocked down A431 cells that migrated during 72h. For each condition, 4 to 5 pictures per experiment of 3 independent experiments were analysed. **(E)** Representative bright-field images of the gaps between control and p120-catenin knocked down A431 cells sheets after 0, 24, 48 and 72h of migration. The yellow lines represent the contour of the gaps not covered by the cells (wound area). Scale bar = 200 µm. **(F)** Wound area after 0, 24, 48 and 72h of migration of control and p120-catenin knocked down A431 cells. For each condition, 4 pictures per experiment obtained in 3 independent experiments were analysed. All the data are presented as mean ± s.d. and tested by Two-way ANOVA (ns: not significant; **: p<0,01; ****: p<0,0001)

Since EGTA is non-specific towards adherens junctions (Clapham, 2007), we generated A431 cells knocked down for p120-catenin. This protein is known to stabilize cadherins at the plasma membrane by binding to their juxtamembrane domain (Ishiyama et al., 2010) and its depletion typically results in cell individualization (Davis et al., 2003, Macpherson et al., 2007). The efficiency of p120-catenin knockdown using short-hairpin RNA (shRNA) was confirmed by Western-Blot and immunofluorescence, with a significantly reduced p120 expression compared to cells treated with a non-target shRNA control (Supplementary Fig. 2B-D).

We evaluated the effect of the p120-catenin knockdown. As expected, it led to the individualization of many A431 cells at the sheet margin compared to the control condition (Fig. 6B). This was coupled with a loss of the supracellular actin network in these areas, as evidenced by a very significant decrease in the number of hubs and the total length of cables (Fig. 6B-D). To demonstrate that the effect on ASAN was not solely due to a physical separation of the cells, we looked at the apical actin organization at the rear of migration where cells remained constrained (Fig. 6B). We observed a similar decrease in the number of hubs and total length of cables in this area (Fig. 6B-D). This demonstrates that cadherin-based adherens junctions are necessary for the formation of ASAN in A431 cells.

Interestingly, p120-catenin knockdown cells still migrated and took significantly less time to close the gap between both cell sheets compared to control cells (Fig. 6E and F). Moreover, we did not observe fingers (Supplementary Fig. 2E), likely due to the individualization of cells at the migration front. These results show that cadherin-based adherens junctions are also essential for the establishment of fingers during A431 cell migration.

To explore whether other junctions contribute to ASAN’s formation, we knocked-down Zonula Occludens 1 (ZO-1), a protein belonging to the tight junctions (Zihni et al., 2016) (Supplementary Fig. 2F). We found that ZO-1 knockdown had no effect on the supracellular actin network, with the number of hubs, their area and the length of cables remaining the same than in control cells (Supplementary Fig. 2G-J). Overall, our data indicate that adherens junctions, but not tight junctions, are required for the formation of the supracellular actin network in A431 cells.

### ROCK-mediated contractility is required for the formation of ASAN in A431 cells

Many supracellular actin structures are known to contain phosphorylated non-muscle myosin II (NMII) (Köppen et al., 2006, Tamada et al., 2007, Röper, 2012, Reffay et al., 2014, Wang et al., 2020). Phosphorylation of the NMII regulatory light chain promotes NMII filament assembly and increases its ATPase activity, enabling NMII to slide anti-parallel actin filaments along each other, producing contractile forces (Murrell et al., 2015). We hypothesized that ASAN might be contractile in A431 cells. To investigate this, we examined the localization of the phosphorylated form of myosin light chain 2 (pMLC2). We observed that many hubs and cables stained for pMLC2, indicating that ASAN in A431 cells is partially contractile (Fig. 7A).

**Figure 7.**
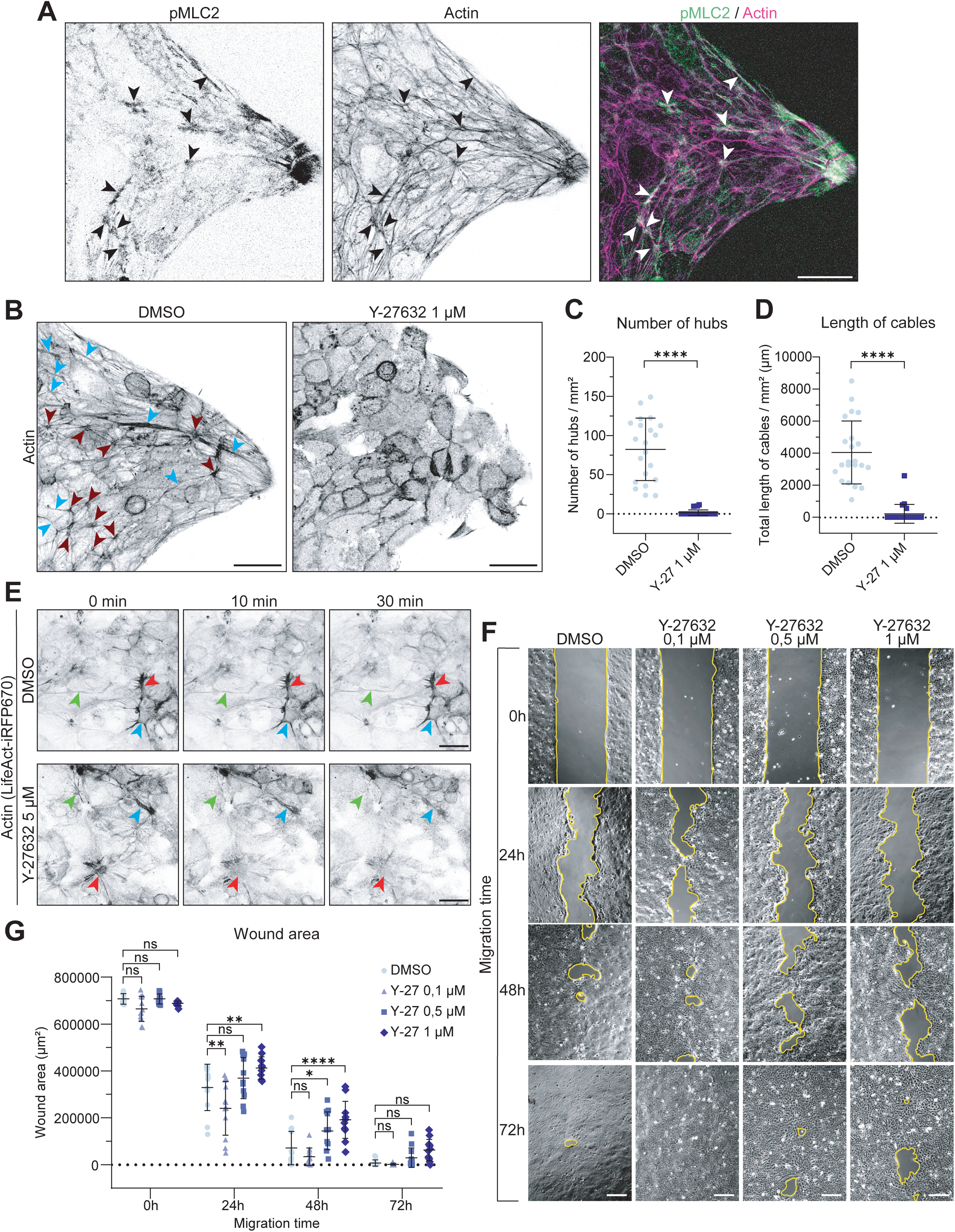
Contractility mediated by ROCK is essential for the formation and maintenance of ASAN and the efficient collective migration of A431 cells. **(A)** Representative maximum intensity projections of the apical Z-planes of a finger formed after 72h of A431 migration. The phospho-myosin light chain 2 (pMLC2) and actin were stained. Black and white arrowheads point towards supracellular actin structures that colocalize with the pMLC2 staining. Scale bar = 50 µm. **(B)** Representative maximum intensity projections of the apical Z-planes of fingers formed after 72h of migration of A431 treated with DMSO (control) or 1 µM of the ROCK inhibitor Y-27632. Cells were stained for actin. Red and blue arrowheads point towards hubs and cables. Scale bar = 50 µm. **(C-D)** Number of hubs per mm² (C) and total length of cables per mm² (D) after 72h of treatment of A431 cells with DMSO (control) or 1 µM of Y-27632. For each condition, 7 to 8 pictures per experiment of 3 independent experiments were analysed. **(E)** Representative maximum intensity projections of the apical Z-planes of a part of an A431 cell sheet that migrated during 72h before being treated with DMSO (control, top) or 5 µM of Y-27632 (bottom). 0 min corresponds to the beginning of the treatment. Cells are stably expressing the actin probe LifeAct-iRFP670. Green, red and blue arrowheads point towards cables and hubs that are maintained in the control condition but disappear after Y-27632 treatment. Scale bar = 50 µm. **(F)** Representative bright-field images of the gaps between sheets of A431 cells treated with DMSO (control) or different concentrations of Y-27632 after 0, 24, 48 and 72h of migration. The yellow lines represent the contour of the gaps not covered by the cells (wound area). Scale bar = 200 µm. **(G)** Wound area after 0, 24, 48 and 72h of migration of A431 cells treated with DMSO (control) or different concentrations of Y-27632. For each condition, 4 pictures per experiment obtained in 3 independent experiments were analysed. All the data are presented as mean ± s.d. and tested by Mann-Whitney (C, D) or Two-way ANOVA (G) (ns: not significant; *: p<0,05; **: p<0,01; ****: p<0,0001)

Different pathways can promote contractility through myosin light chain phosphorylation. One key player is the Rho-associated kinase (ROCK), a well-known effector of the Rho GTPase (Matsui et al., 1996), that induces contractility either directly by phosphorylating and activating the myosin light chain (Amano et al., 1996) or indirectly by phosphorylating and inactivating the myosin light chain phosphatase (Kimura et al., 1996). To investigate the role of ROCK-mediated contractility in the formation of ASAN, we treated A431 cells with Y-27632, a ROCK inhibitor (Ishizaki et al., 2000), from the onset of migration and during 72 h. This treatment resulted in a complete disappearance of the supracellular actin network (Fig. 7B), evidenced by a significant decrease in the number of hubs and total length of cables (Fig. 7C and D). Importantly, after washing out the Y-27632 treatment, some cables and hubs reappeared within a few hours, demonstrating the reversibility of the process (Supplementary Fig. 3A and Movie 4). These findings demonstrate that the contractility mediated by ROCK is required for ASAN formation.

Time-lapse imaging of cells stably expressing a fluorescent actin probe revealed a fast effect of a 5 µM Y-27632 treatment applied after 72 h of migration (Fig. 7E and Movies 5 and 6). Indeed, ASAN completely disassembled within 30 min of treatment. This result indicates that ROCK-mediated contractility is not only required for the formation, but also for the maintenance the supracellular actin network in A431 cells, potentially through the actin crosslinking properties of NMII (Vicente-Manzanares et al., 2009).

We next assessed how ROCK inhibition affected the overall migration process. Interestingly, Y-27632 produced two different responses: the lower concentration of the inhibitor significantly increased wound closure, while higher concentrations significantly reduced it (Fig. 7F-G). This dual effect of ROCK inhibition suggests that a precise balance of contractility is required for optimal collective migration at a specific speed. Interestingly, despite the complete loss of ASAN after 72 h of Y-27632 treatment, there were no significant changes in the number and area of fingers. Only the highest concentration of Y-27632 resulted in a small but significant increase in finger elongation (Supplementary Fig. 3B-E). This may be attributed to the persistence of some actin arcs post-treatment which may still partially restrict protrusion formation in the side cells at the migration front. Thus, while ROCK inhibition disrupts ASAN, it primarily affects the speed of wound closure but has minimal impact on finger formation. This suggests that ASAN might not be essential for finger formation. However, it remains unclear whether the disappearance of ASAN is directly linked to the changes in cell migration speed.

### MAP4K4 inhibition enhances the density of ASAN in A431 cells

Previous work from our lab demonstrated that the mitogen-activated kinase kinase kinase kinase 4 (MAP4K4), by being localized at adherens junctions, promotes their disassembly in small clusters of A431 cells (Alberici Delsin et al., 2023). This primarily occurs through a reorganization of the actomyosin cytoskeleton and a decrease in tension loading at these junctions (Alberici Delsin et al., 2023). Specifically, MAP4K4 loss-of-function resulted in an increase in contractile dorsal stress fibers and adherens junction stability through an increase in tension at these junctions. Due to these findings, we used MAP4K4 loss-of-function as a tool to stabilize adherens junctions and investigate its effects on the supracellular actin network. We inhibited MAP4K4 using GNE-495. After 72 h of treatment, the supracellular actin network remained present but exhibited an increased density compared to the control condition (Fig. 8A). This was evidenced by a significant increase in the number of hubs and total length of cables while the area of hubs remained unchanged (Fig. 8B-D). These results suggest that stabilizing adherens junctions and increasing tension on these junctions promote the formation of denser apical supracellular actin structures.

**Figure 8.**
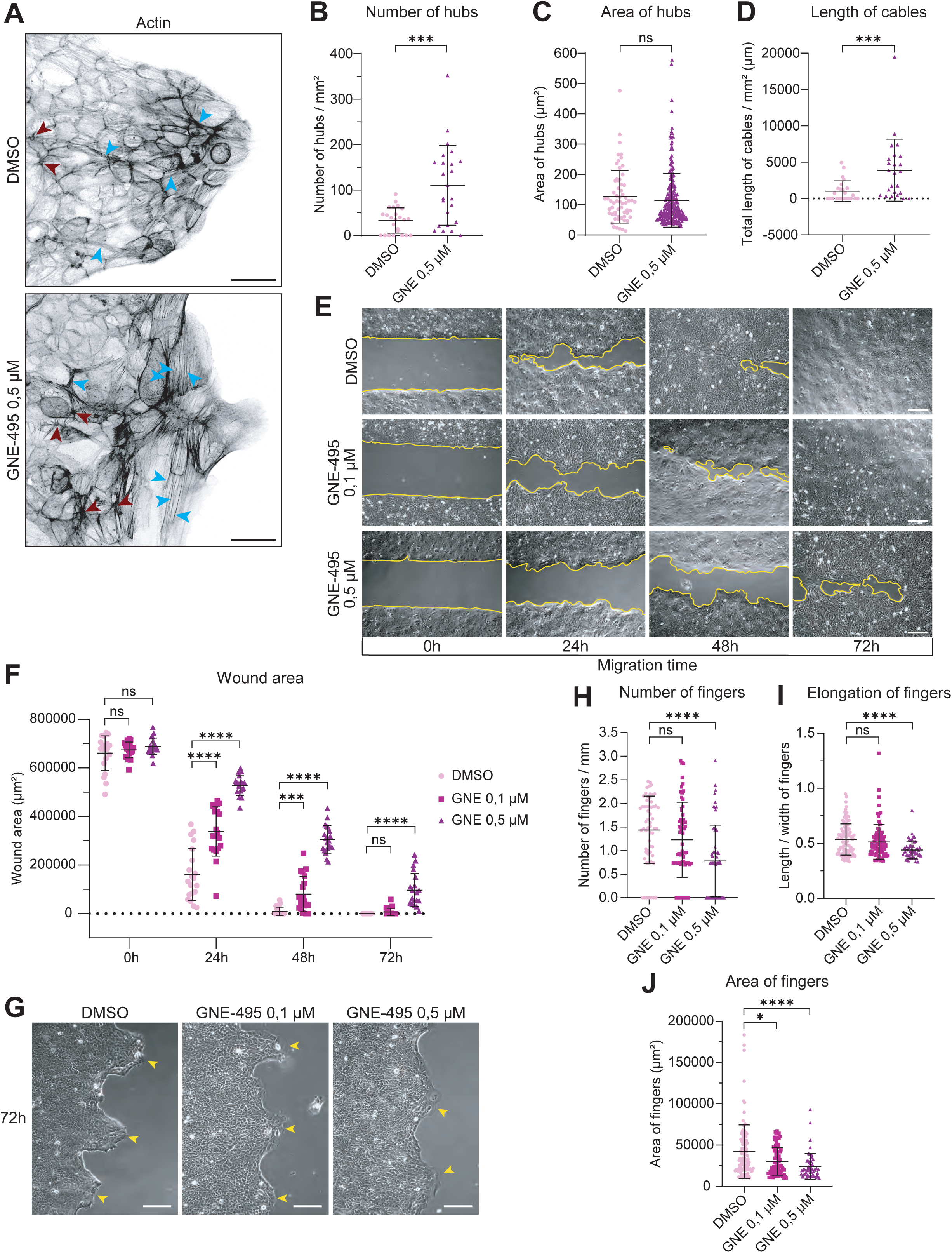
MAP4K4 inhibition enhances the density of ASAN and impedes the collective migration of A431 cells. **(A)** Representative maximum intensity projections of the apical Z-planes of fingers formed after 72h of migration of A431 treated with DMSO (control) or 0,5 µM of the MAP4K4 inhibitor GNE-495. Cells were stained for actin. Red and blue arrowheads point towards hubs and cables. ––Scale bar = 50 µm. **(B-D)** Number of hubs per mm² (B), area of hubs (C) and total length of cables per mm² (D) after 72h of treatment of A431 cells with DMSO (control) or 0,5 µM of GNE-495. For each condition, 8 pictures per experiment obtained in 3 independent experiments were analysed. **(E)** Representative bright-field images of the gaps between sheets of A431 cells treated with DMSO (control) or different concentrations of GNE-495 after 0, 24, 48 and 72h of migration. The yellow lines represent the contour of the gaps not covered by the cells (wound area). Scale bar = 200 µm. **(F)** Wound area after 0, 24, 48 and 72h of migration of A431 cells treated with DMSO (control) or different concentrations of GNE-495. For each condition, 4 pictures per experiment obtained in 5 independent experiments were analysed. **(G)** Representative bright-field images of the periphery of sheets of A431 cells after 72h of migration and treatment with DMSO or different concentrations of GNE-495. The yellow arrowheads point towards fingers. Scale bar = 200 µm. **(H-J)** Mean number of fingers/mm (H), elongation of fingers (I) and area of fingers (J) formed after 72h of migration of A431 cells treated with DMSO or different concentrations of GNE-495. The elongation was calculated by the ratio length/width of fingers and the area by the entire surface over the base of the fingers. For each condition, 12 pictures per experiment obtained in 5 independent experiments were analysed. All the data are presented as mean ± s.d. and tested by Mann-Whitney (B-D), Two-way ANOVA (F) or Kruskal-Wallis (H-J) (ns: not significant; *: p<0,05; ***: p<0,001; ****: p<0,0001)

To assess the kinetics of these changes, we let A431 cells migrate for 72 hours without the inhibitor and treated them with GNE-495 for short periods of time. Interestingly, only 6 h of treatment were sufficient to induce a significant increase in the number of hubs and the total length of cables (Supplementary Fig. 4A, B and D). However, unlike the long treatment, short MAP4K4 inhibition led to a significant decrease in the area of hubs (Supplementary Fig. 4C). This suggests that newly formed hubs induced by MAP4K4 inhibition may enlarge over time until they reach a regular size.

When looking at the effect of three different concentrations of GNE-495 on A431 collective migration, we observed that wound closure was significantly slowed in a dose-dependent manner (Fig. 8E and F). Importantly, MAP4K4 inhibition with the highest concentration of GNE-495 also led to a significant decrease in finger number, elongation and area (Fig. 8G-J). Thus, MAP4K4 inhibition enhances the density of ASAN and reduces cell migration. Further investigation could reveal whether there is a direct link between ASAN’s density and the speed of migration.

### Contractile ASAN is a source of tension and long-range force-transmission at the apical side of A431 cells

Our previous findings demonstrated that the supracellular actin network is a contractile structure (Fig. 7A). We next examined whether ASAN is under tension. To test this, we performed ablation experiments using a two-photon laser on A431 cells stably expressing a fluorescent actin probe (LifeAct-iRFP670) and junctional marker (E-cadherin-mCitrine). Severing specific cables or hubs resulted in a recoil of these actin structures on both sides of the cut, confirming that the network is indeed under tension (Fig. 9A, Movie 7).

**Figure 9.**
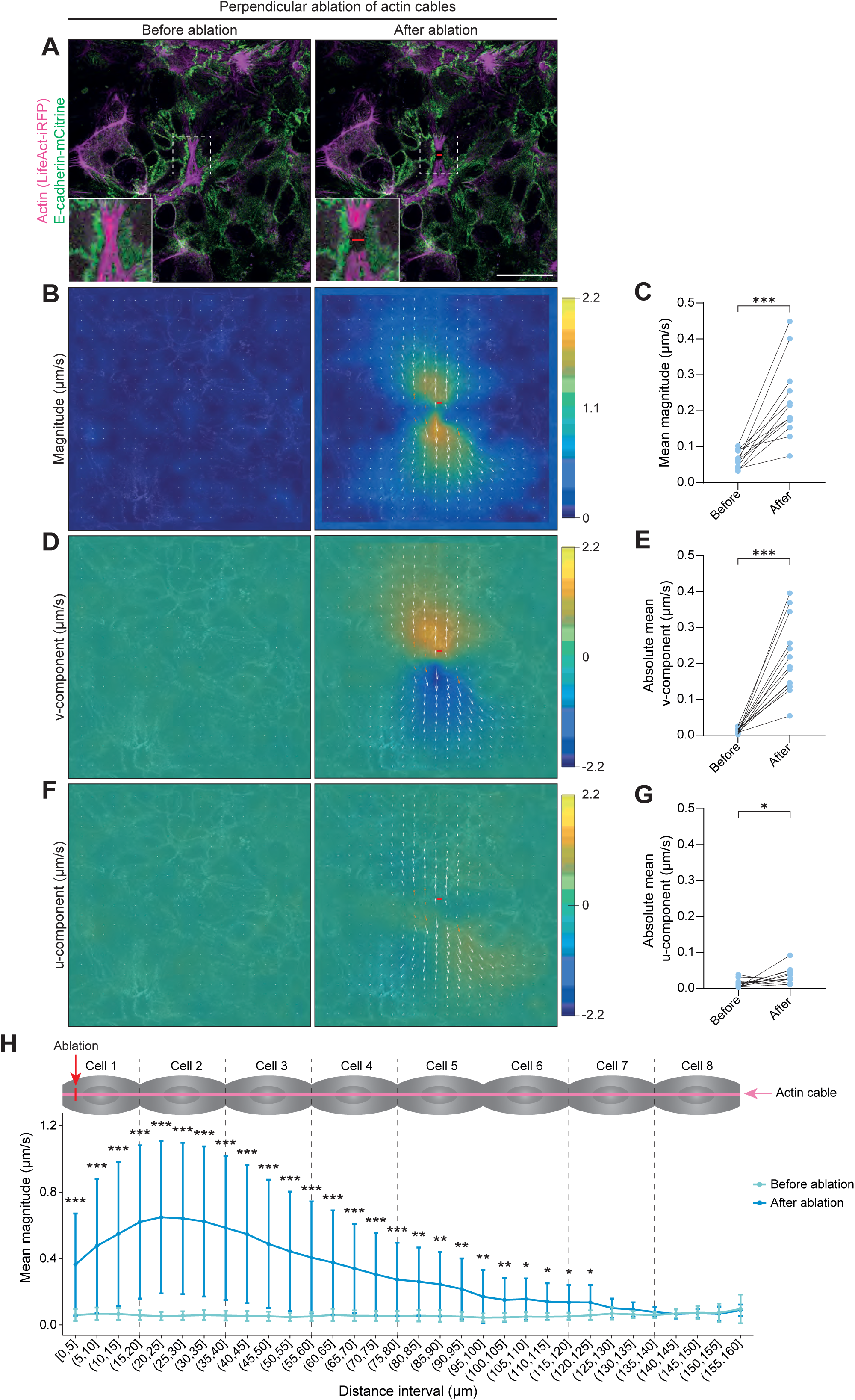
ASAN transmits tension at a multicellular level in A431 cells. **(A)** Confocal images of the apical part of an A431 cell sheet after 72 h of migration, before and after perpendicular laser ablation of an actin cable. Cells stably express the actin probe LifeAct-iRFP670 and the adherens junction protein E-cadherin-mCitrine. Red lines indicate the ablation site. Images were acquired at 1 sec interval. Insets show magnified views of the regions outlined by dotted squares. Scale bar = 50 µm. **(B, D, F)** Colormaps representing the magnitude (B), vertical (v-component, D) and horizontal (u-component, F) movement of junctions labeled with E-cadherin-mCitrine, before and after ablation perpendicular ablation of the actin cable shown in (A). The color scales (right) indicate displacement intensity for each parameter. White arrows represent the direction and magnitude of the movement at each point. Red lines indicate the ablation site. **(C, E, G)** Quantification of junctional movements marked by E-cadherin-mCitrine following perpendicular ablation of actin cables. Shown are the mean magnitude (C), vertical (v-component, E) and horizontal (u-component, G) displacement of vectors on each side of the cut. For each parameter, 13 movies obtained in 3 independent experiments were analysed. **(H)** Magnitude before and after perpendicular ablation of actin cables, plotted as a function of distance from the severed site. Significant membrane displacement is observed up to 7 cells. 13 movies obtained in 3 independent experiments were analysed. Data are presented as individual data points (C, E, G) or mean ± s.d. (H) and tested by Wilcoxon (C, E, G) or t-test (H) (*: p<0,05; **: p<0,01; ***: p<0,001).

Using Particle Image Velocimetry (PIV) analysis on the Ecadherin-mCitrine signal, we were able to follow adherens junctions displacements before and after ablation of supracellular actin cables (Fig. 9). The mean magnitude of junctional movement increased significantly following ablation (Fig. 9B, C and H), showing that severing a supracellular actin cable leads to significant tension release in neighboring cells. As a control, we severed apical cortical actin structures that do not belong to ASAN (Supplemental Fig. 5A, Movie 8). The mean magnitude of junction displacement post-ablation was markedly lower, as it did not exceed 0.1μm/s (Supplemental Fig. 5B), compared to over 0.4μm/s following ablation of supracellular cables (Fig. 9C). Consistently, the u- and v-component did not change much after ablation of cortical actin (Supplementary Fig. 5C and D). This supports the conclusion that ASAN is subjected to considerably higher mechanical tension than cortical actin structures.

To ensure consistency in PIV measurements, all severed actin cables were oriented vertically in the frame. Measurements of the v-component (vertical displacement in the y-axis) and u-component (horizontal displacement in the x-axis) of junctions’ movement revealed that displacement after ablation of supracellular actin cables occurred predominantly along the y-axis with minimal displacement along the x-axis (Fig. 9.D-G, Movie 9). This indicate that tension release primarily occurred along the axis of the severed cable.

To assess whether the directionality of tension release depends on the orientation of ablation, we also performed ablations with the cut oriented in the axis of the cable (Supplementary Fig. 5E). These ablations induced a significant increase in displacement magnitude (Supplementary Fig. 5F), comparable to when the cut was oriented perpendicularly to supracellular actin cables (Fig. 9B and C). For parallel ablations of supracellular actin cables, tension was also mainly released in the axis of the ablated cable, with minimal movement perpendicular to it (Supplementary Fig. 5G-H) This suggests that the direction of tension release is independent of the orientation of ablation, but solely depends on the orientation of the severed cable.

Further analysis revealed that tension release upon supracellular cable ablation propagates over long distances (Fig. 9H). While the largest displacements were observed in the first two adjacent cells, measurable movement was still detected up to approximately 125 µm away, equivalent to about seven cells away. These findings suggest that, beyond bearing tension, ASAN may function as a long-range force transmission device, capable of propagating tensile forces across the epithelial sheet.

## DISCUSSION

In this study, we describe an apical supracellular actin network (ASAN) present in skin epithelial cells. This network is composed of interconnected star-shaped hubs and long cables that extend across multiple cells through adherens junctions. It is contractile and applies tension on the apical surface of A431 skin squamous carcinoma cells. Furthermore, it can transmit forces over long distances. During migration, it densifies, as it exhibits more supracellular actin cables and hubs as the cells migrate and aligns with the direction of migration. Overall, our data suggest that ASAN may facilitate mechanical communication within sheets of epidermal cells through long-range force transmission.

Our finding originated from a screening of various cell lines that revealed that HaCaT epidermal keratinocytes, A431 and HSC-5 skin squamous carcinoma cells, and CAL27 tongue squamous carcinoma cells migrate collectively by forming multicellular hierarchical structures called fingers. These fingers, with varying prominence across cell lines, feature large protruding cells at their tips followed by strands of cells of different widths and lengths. Already observed in collectively migrating MDCK canine kidney epithelial cells (Haga et al., 2005, Poujade et al., 2007, Petitjean et al., 2010, Reffay et al., 2011, Reffay et al., 2014, Yamaguchi et al., 2015, Balcioglu et al., 2020, Isozaki et al., 2020) and IAR-2 rat liver epithelial cells (Omelchenko et al., 2003), these multicellular structures could be viewed as the 2D equivalent of branches seen during the collective invasion of squamous carcinoma cells in spheroids and tumour specimens (Dissanayaka et al., 2012, Kato et al., 2023). Consistent with earlier observations in scratch assays (Goel et al., 2012, Pedrazza et al., 2020), U2OS osteosarcoma cells and A549 lung adenocarcinoma cells did not form fingers. This propensity of normal and cancerous epithelial cells to form fingers across various species suggests a conserved mechanism of collective migration in epithelial tissues.

Given the role of supracellular actin structures in collective cell migration across many models, we investigated the actin organization in epithelial cell lines during their collective movement. It revealed that epithelial cells forming fingers such as A431 cells exhibited two types of supracellular structures: 1) basal arcs at the lateral side of fingers, and at the neck connecting fingers to the rest of the epithelial sheet, and 2) a dense, apical supracellular actin network (ASAN). Other cell lines tested were devoid of the arcs and of ASAN, suggesting that squamous epithelial cells are more prone to form supracellular actin structures compared to cell lines from different origins. The basal arcs resemble structures observed in MDCK cells that were shown to be under tension, restricting protrusion formation and transmitting forces between leading-edge cells on finger sides to facilitate their coordinated movement (Reffay et al., 2014). Consistent with this, we did not see protrusions in cells containing these actin arcs.

We focused our study on ASAN and found that it localized near the apical plasma membrane and that it is composed of star-shaped hubs interconnected with long cables spanning multiple cells. Similar structures have been observed in other contexts. For instance, ventral furrow cells in Drosophila show an apical network of interconnected hubs that promotes the apical cell constriction (Martin et al., 2009). Such networks of hubs have also been described in the basal epithelium of the Ephydatia mülleri sponge (De Ceccatty, 1986). Regarding the cables, many models showed the presence of a basal supracellular actin belt at the leading-edge of migration, oriented perpendicular to the direction of movement and acting as a contractile purse-string (Bement et al., 1993, Kiehart et al., 2000, Panfilio and Roth, 2010, Czerniak et al., 2016). Similar actomyosin cables at the apical side of various tissues during embryonic development promote cell invagination for processes like salivary gland formation and neural tube closure (van Straaten et al., 2002, Röper, 2012). Additionally, long actin cables have been described to segregate different cell types or compartments in Drosophila, such as those delimitating rhombomeres (Monier et al., 2010). Unlike the supracellular actin structures formed at the leading-edge migration front, the network of actin hubs and cables in A431 cells is located within the cell sheet. Interestingly, mouse skin keratinocytes have apical actin cables that seem to be continuous between neighbour cells and interconnected at junctions (Vaezi et al., 2002). Visually, the apical actin network in skin keratinocytes seem very similar to what we observed in HSC-5 cells although its span is limited to pairs of neighbouring cells.

Our data indicate that the supracellular network requires adherens junctions to form. Indeed, we found that the disassembly of adherens junctions by different means leads to the disappearance of ASAN, while stabilization of adherens junctions by inhibiting the kinase MAP4K4 increases the density of ASAN. On the opposite, tight junctions seem dispensable as depletion of ZO-1 does not destabilize the network. The requirement of adherens junctions and the observation that connection between actin bundles overlap with the β-catenin strongly suggest that ASAN is assembled by the cadherin-catenin complex (Buckley et al., 2014).

Furthermore, we found that parts of the actin supracellular network contain the actin bundling protein α-actinin-1 and active non-muscle myosin II, as detected by immunostaining. Previous studies have demonstrated the role of α-actinin-1 in actin filament bundling in ventral, transversal and dorsal stress fibers at the cellular level (Feng et al., 2013, Kemp and Brieher, 2018). At the multicellular scale, the supracellular purse-string actin ring in Drosophila dorsal closure is enriched in α-actinin at the level of the junctions (Rodriguez-Diaz et al., 2008). Hence, we suppose that α-actinin-1 is important to assemble actin into large bundles and into hubs, but this remains to be precisely tested. It is surprising, however, that only parts of ASAN were labelled for α-actinin-1. Although it is possible that anti-α-actinin-1 antibodies cannot penetrate dense structures, our data suggest that some parts of ASAN can subsist without α-actinin-1 or that other actin-bundling proteins might replace α-actinin-1 in the rest of the network. Similarly, active non-muscle myosin II, as detected by pMLC2 staining was present on a subset of ASAN, suggesting that parts of the network are contractile. Treatment with the ROCK inhibitor Y-27632 induced the reversible disassembly of the cables, suggesting that active myosin II is required to maintain ASAN. Our data on myosin II can be explained by three different, non-exclusive mechanisms: 1) contractility applied to the network is required for its maintenance, 2) myosin II crosslinks actin cables with its anti-parallel organisation, in a contractility independent mechanism (Vicente-Manzanares et al., 2009) and 3) contractility frequently stabilizes adherens junction (Thomas et al., 2013, Engl et al., 2014, Chu et al., 2004), hence the reduction of contractility may destabilize cell-cell junctions in A431 cells, as we previously reported (Alberici Delsin et al., 2023).

Actomyosin forces were shown to be transmitted through adherens junctions in previous studies, mostly when actin fibers are perpendicular to the junction (Chen et al., 2018, Yonemura, 2017, Fernandez-Gonzalez and Peifer, 2022). Here, we found that the bundles that form the actin network are distributed perpendicularly to the junctions, suggesting that they apply forces to those junctions. Accordingly, we found that the junctions adopt an interdigitated morphology, indicating that tension is applied to the junction. We also found that vinculin, a protein recruited at adherens junctions when forces are applied on α-catenin, is almost uniformly present at apical cell-cell junctions of A431 cells. We concluded from these observations that the apical surface of A431 cell sheets is under tension, but further studies using more quantitative mechanosensitive probes will be necessary to determine whether mechanical forces are specifically increased at junctions that interconnect the supracellular actin network. To investigate whether ASAN transmits forces directionally, we performed laser ablation of actin cables. In every instance where supracellular actin structures were severed, we observed a recoil. As expected if forces are transmitted by cables, particle image velocimetry analysis of E-cadherin movement revealed that the release of forces was directional, along the axis of the severed cable, with minimal movement perpendicular to it. More strikingly, significant displacements were detected up to 125 µm away from the cut site in both directions. On average, each ablated cable appeared to transmit tension across approximately 12 to 14 cells, *in toto*. These results support the idea that ASAN is a directional, long-range force transmission device.

Our observation that ASAN aligns with the direction of migration suggests that it may play a role in collective cell migration. Accordingly, we found that there is an inverse correlation between the density of ASAN and the migration speed, as treatment with the MAP4K4 inhibitor GNE-495 slows migration while densifying the network, and low doses of the ROCK inhibitor Y-27632 resulted in reduced network, but increased cell velocity. Further experiments would be necessary to directly link the density of ASAN with migration speed as the conditions we used to manipulate the network affects other migration determinants. The network may play other biomechanical functions. Indeed, we found such a network in cells derived from squamous epithelia. This dense ASAN might thus have a specialized function in particular tissues such as the skin or the tongue epithelia that need to have a certain elasticity and be able to heal rapidly when damaged. Furthermore, it would be interesting to investigate whether proteins specifically expressed in the skin, such as cytokeratins or other intermediate filaments can be responsible for the formation of the network.

## MATERIALS AND METHODS

### Cell culture

Human skin squamous carcinoma A431 cell line (#CRL-1555), human tongue squamous carcinoma CAL27 cell line (#CRL-2095) and immortalized human embryonic kidney HEK-293T cell line (#CRL-3216) were purchased from ATCC. Spontaneously transformed human keratinocyte HaCaT cell line (#T0020001) was obtained from Clini Sciences and human skin squamous carcinoma HSC-5 cell line (#JCRB1016) from JCRB Cell Bank. Human lung adenocarcinoma A549 cell line (#CCL-185) and human osteosarcoma U2OS cell line (#HTB-96) were originally purchased from ATCC and were kindly provided by Dr. Sébastien Carréno and Dr. Marc Therrien (IRIC, Université de Montréal, Canada), respectively.

A431, CAL27, HaCaT, A549, U2OS and HEK-293T cells were grown in Dulbecco’s Modified Eagle Medium (DMEM; Gibco; #11995073) supplemented with 10 % fetal bovine serum (FBS; Wisent; #080-150) and 1 % penicillin-streptomycin (Gibco; #15140122). HSC-5 cells were cultured in Iscove’s Modified Dulbecco’s Medium (IMDM; Gibco; #12440053) supplemented with 10 % FBS and 1 % penicillin-streptomycin. All cell lines were maintained in a humidified atmosphere at 37 °C with 5 % CO_2_ and were periodically tested for mycoplasma contamination using PCR.

A431, CAL27 and HSC5 cells were plated at a density of 3500-4500 cells/cm² and passed twice a week upon reaching cluster confluence. HaCaT cells, were plated at a higher density of 7000 cells/cm² and were passed twice a week once they reached 40-50 % confluence. A549 cells were plated at 5000 cells/cm² and passed twice a week once they reach 80-90 % confluence. U2OS cells, plated at a density of 12000 cells/cm², were also passed twice a week at 80-90 % confluence. HEK-293T cells were plated at a density of 45000-90000 cells/cm² and passed three times a week when they reach 80-90 % confluence.

### Plasmids

All plasmids were amplified in One Shot Stbl3 Chemically Competent Escherichia coli (Invitrogen; #C737303) and purified using Qiagen Plasmid Midi kit (#12143).

Lentiviral packaging plasmid psPAX2 (Addgene; plasmid # 12260) and envelope plasmid pCMV-VSV-G (Addgene; plasmid # 8454) were gifts from Didier Trono and Bob Weinberg, respectively. To produce lentiviruses, these were used in conjunction with different transfer plasmids presented here after.

pLenti LifeAct-EGFP BlastR (Addgene; plasmid # 84383) and pLenti Lifeact-iRFP670 BlastR (Addgene; plasmid # 84385), used to express the LifeAct filamentous actin probe, were given by Ghassan Mouneimn. A lentiviral plasmid for the expression of the fluorescent adherens junction protein E-cadherin-mCitrine was kindly shared by Arnold Hayer (McGill University, Montréal, Canada). pLKO.1 plasmids containing shRNA came from the MISSION shRNA plasmid DNA library and were provided by IRIC’s High-Throughput Core Facility (Université de Montréal, Canada). The following shRNA constructs were used: CTNND1 shRNA (Sigma-Aldrich; #TRCN0000122986; target sequence: 5’-CGGATATACATCTCACTTCTT-3’) to target the adherens junction protein p120-catenin, TJP1 shRNA (Sigma-Aldrich; #TRCN0000116252; target sequence: 5’-GCAATAAAGCAGCGTTTCTAT-3’) to target the tight junction protein ZO-1 and non-target shRNA (Sigma-Aldrich; #SHC002; target sequence: 5’-CAACAAGATGAAGAGCACCAA-3’) as a control that targets no known mammalian genes.

### Production of lentiviruses and infections of cells

Lentiviruses were produced by first seeding 5 × 10⁶ HEK-293T cells in 10 cm culture dishes containing 10 mL of complete culture medium. The next day, 6 µg of transfer plasmid, 3 µg of psPAX2 packaging plasmid, 3 µg of pCMV-VSV-G envelope plasmid and 36 µL of 1 mg/mL polyethylenimine (PEI; Polysciences; #23966) were vortex-mixed in Opti-MEM (Gibco; #31985070) to a final volume of 1200 µL. This solution was incubated for 15 min at room temperature before being added droplet-wise to the HEK-293T. After 16 h, the transfection medium was replaced by 7 mL of fresh complete growth medium. Medium containing lentiviral particles was collected 48 h after transfection, filtered through 0,45 µm filters (Sarstedt; #83.1826) and mixed with 200 µL of Hepes 1 M pH 7 (Bioshop; #HEP001.50) before being stored at -80 °C.

For lentivirus infection, 3 × 10^5^ A431 cells were seeded in 10 cm culture dishes containing 10 mL of complete growth medium. The next day, this medium was replaced with 5 mL of fresh complete medium. 3 mL of the virus-containing medium was mixed with 15 µL of 4 mg/mL polybrene (Sigma-Aldrich; #H9268-10G) and added to the A431 cells. Selection of lentivirus-transduced cells started 48h after infection with 0,75 µg/mL puromycin (InvivoGen; #ANT-PR) and/or 1 µg/mL blasticidin (InvivoGen; #ANT-BL).

### Collective cell migration assay

For immunofluorescence experiments, 2-well culture inserts (IBIDI, #80209) were fixed directly onto glass coverslips. For time-lapse imaging and plasma membrane observations, inserts were placed into 2-well glass-bottom µ-slides (IBIDI; #80287). 100 µL of complete growth medium containing a certain number of cells were added into each well of the insert. The following cell numbers per well were used for each cell line: 1 × 10⁵ A431 cells, 7,5 × 10^4^ CAL27 cells, 2,5 × 10^4^ HSC-5 cells, 6 × 10^4^ HaCaT cells, 5 × 10^4^ A549 cells or 4 × 10^4^ U2OS cells. After 24 h of incubation, cells reached 100 % confluence. The inserts were then gently removed using tweezers and cells were rinsed three times with complete growth medium. This was designated as the 0 h time point of migration. Cell migration into complete growth medium, with or without drug treatment, was monitored for up to 72 h (3 days).

To determine individual cell contributions to the actin network, a mosaic of A431 cells was generated by mixing an equal number of LifeAct-EGFP expressing cells and LifeAct-iRFP670 expressing cells.

### Treatment with inhibitors and EGTA

The calcium chelator EGTA (Bioshop; #EGT101.25) was solubilized in distilled water at a 50 mM concentration and filtered with a 0,2 µm filter (Sarstedt; #83.1826.001). This stock solution was stored at 4 °C. Stock solutions of the ROCK inhibitor Y-27632 (New England Biolabs; #13624S) and the MAP4K4 inhibitor GNE-495 (MedChemExpress; #HY-100343) were prepared by dissolving the powders into DMSO (Bioshop; #DMS666) to a 10 mM concentration and stored at -20 °C and -80 °C, respectively. These stock solutions were then diluted at different concentrations to test on the cells.

#### Live imaging to follow actin after adherens junction destabilization

The effect of adherens junction destabilization on the supracellular actin network was monitored by live imaging after treating migrating A431 cells with 4mM EGTA or distilled water (control). A431 cells stably expressing LifeAct-iRFP670 and E-cadherin-mCitrine were seeded in a 2 well glass-bottom µ-slide (IBIDI; #80287) as described in the “Migration assay” section. After 72 h of migration, the regular DMEM-based complete growth medium was replaced with 1 mL of a live-imaging compatible medium (FluoroBrite DMEM (Gibco, #A1896701) supplemented with 580 mg/L L-glutamine (Gibco; #25030149), 110 mg/L sodium pyruvate (Gibco; #11360070), 10 % FBS (Wisent; #080-150) and 1 % penicillin-streptomycin (Gibco; #15140122)). EGTA was diluted to 8 mM in the live-imaging compatible medium. Equivalent volume of distilled water was added in the control. After 10 min of acquisition on a confocal microscope, and without stopping the acquisition, 1 mL of the 8 mM EGTA solution (or diluted distilled water for control) was injected to achieve a final concentration of 4 mM. Conditions of image acquisition are specified in the “Image acquisition” section.

#### Live imaging to follow actin after ROCK inhibition

The effect of ROCK inhibition was assessed following the same procedure as above, except that cells were treated with 5 µM Y-27632 or DMSO (control). Y-27632 was diluted to 10 µM in the live imaging compatible medium. DMSO was diluted similarly for the control.

#### Live imaging to follow the recovery of the supracellular actin network

To observe the reappearance of the supracellular actin network after removing EGTA or Y-27632, A431 cells stably expressing LifeAct-iRFP670 and E-cadherin-mCitrine were seeded in a 2 well glass-bottom µ-slide (IBIDI; #80287) as described in the “Migration assay” section. After 72h of migration, cells were treated with 4 mM EGTA for 20 min or 5 µM Y-27632 for 40 min. EGTA and Y-27632 were diluted into complete growth medium. Cells were then washed three times with complete growth medium and 1 mL of the live imaging compatible medium described above was added. Image acquisition on a confocal microscope started following conditions described under “Image acquisition”.

#### Long-term ROCK inhibition

To assess the effect of prolonged ROCK inhibition on the collective cell migration and the supracellular actin network, A431 cells were treated with 0,1, 0,5 and 1 µM Y-27632 diluted in complete growth medium for 72 h after removing the silicon inserts. DMSO, diluted similarly to the highest Y-27632 concentration, served as a control. Treatments were renewed daily.

#### MAP4K4 inhibition

The effect of the MAP4K4 inhibition was evaluated by treating A431 cells either with 1 µM GNE-495 for 6 h after 72 h of migration or with 0,1 µM or 0,5 µM for 72h after removing the inserts. Complete growth medium containing the same volume of DMSO as the one used for the highest GNE-495 concentration was used as a control. For the 72h treatment, the GNE-495 containing medium was renewed every day.

### Immunofluorescence staining

Cells on coverslips that migrated during 0, 24, 48 or 72 h were fixed in 4 % paraformaldehyde (PFA; Fisher Scientific; #50-980-487) for 30 min at room temperature (RT). Note that the actin supracellular structures in A431 disappear rapidly when cells are removed from the incubator, hence cells must be fixated immediately after removal from the incubator. Fixed cells were washed three times with 1X phosphate buffered saline (PBS; Bioshop; #PBS404). They were then permeabilized for 3 min in 0,1 % Triton X-100 (Millipore Sigma; #X100) and washed three times with 1X PBS. Blocking was performed with 2 % bovine serum albumin (BSA; Bioshop; #ALB001) for 30 min to 2 h at RT.

Primary antibodies diluted in 2 % BSA were incubated for 2-3 h at 37 °C or overnight (ON) at 4 °C, depending on the antibody. The following primary antibodies were used : anti-α-actinin 1 rabbit polyclonal antibody (1:250; ON at 4 °C; US Biological; #A0761-02F); anti-β-catenin mouse monoclonal antibody (1:100; 2 h at 37 °C or ON at 4 °C; BD Biosciences; #610153); anti-fibronectin rabbit polyclonal antibody (1:500; ON at 4 °C; Abcam; #ab2413); anti-p120-catenin rabbit polyclonal antibody (1:1000; 2 h at 37 °C; Santa Cruz Biotechnology; #sc-13957); anti-phospho myosin light chain 2 (pMLC2) rabbit polyclonal antibody (1:100; ON at 4 °C; Cell Signaling; #3671) and anti-vimentin rabbit polyclonal antibody (1:500; ON at 4 °C; Abcam; #ab137321). Following this incubation with primary antibodies, cells were washed three times with 2 % BSA.

Secondary antibodies diluted in 2 % BSA were incubated for 1 h at RT, with or without phalloidin and DAPI depending on the experiments. For experiments requiring only actin and/or nuclear staining, cells were permeabilized and washed as described above, then directly incubated with phalloidin and/or DAPI, omitting the antibody steps. The following secondary antibodies were used at 1:1000 dilution: Alexa Fluor 488 goat anti-mouse Immunoglobulin G (IgG) (Invitrogen; #A-11029), Alexa Fluor 488 goat anti-rabbit IgG (Invitrogen; #A-11008), Alexa Fluor 555 goat anti-mouse IgG (Cell Signaling Technology; #4409S) and Alexa Fluor 555 goat anti-rabbit IgG (Cell Signaling Technology; 4413S). For nuclei staining, DAPI (Sigma-Aldrich; #D8417) was used at 1 μg/mL. F-actin was stained using Alexa Fluor 488 phalloidin (1:1000; Invitrogen; #A12379), Alexa Fluor 555 phalloidin (1:750; Invitrogen; #A34055) or Alexa Fluor 647 phalloidin (1:50; Invitrogen; #A22287). After incubation, cells were washed three times with 2 % BSA and once with 1X PBS. Coverslips were mounted on glass slides using Vectashield Plus antifade mounting medium (Vector Laboratories; #H-1900) and sealed using nail polish before image acquisition with a confocal microscope.

To determine the apical localization of the supracellular actin network, plasma membranes were visualized without cell flattening caused by the mounting of coverslips on a slide. Cells were plated in a 2 well glass-bottom µ-slide (IBIDI; #80287). After 72 h of migration, cells were fixed and washed as described above. They were then incubated with 5 µg/mL Wheat Germ Agglutinin Alexa Fluor 488 (Invitrogen, #W11261) diluted in 1X PBS for 10 min at RT. Cells were washed three times with 1X PBS before being permeabilized, blocked and stained as described above. For the final step, Vectashield was added directly to the well in order to cover the cells entirely, and images were acquired immediately after.

### Image acquisition

#### Bright-field microscopy

Bright-field images were captured using a Leica DMRiB inverted microscope equipped with a 10x HC PL Fluotar NA 0.3 Ph 1 objective and a QImaging Retiga EXi digital camera. Image acquisition was performed using the QCapture Suite version 2.9 software.

#### Fluorescence microscopy on fixed cells

Fluorescence microscopy imaging on fixed cells was conducted on a Zeiss LSM880 confocal microscope using Zen software. For vimentin and fibronectin experiments, a 20x Plan-Apochromat NA 0.8 M27 objective was used. All other regular fluorescence microscopy experiments on fixed cells employed a 40x EC Plan-Neofluar NA 1.3 oil immersion objective.

#### High-resolution fluorescence microscopy on fixed cells

High resolution images were acquired on the Zeiss LSM880 confocal microscope using a 63x Plan-Apochromat NA 1.4 DIC M27 oil immersion objective, an Airyscan detector and the Zen software for image acquisition and Airyscan processing. Figure legends indicate when images were acquired using this high-resolution microscopy technique.

#### Time-lapse imaging

A Zeiss LSM700 confocal microscope with a 10x N-Achroplan NA 0.25 Ph1 M27 objective was used to follow protrusion dynamics and analyse cell velocities. Images were acquired every 1.5 min. A Zeiss LSM880 confocal microscope equipped with a 40x EC Plan-Neofluar NA 1.3 oil immersion objective was employed for EGTA and Y-27632 experiments. Post-treatment images were acquired every 2 min, while post-washout images were captured every 5 min. Finally, the laser ablation experiments were done on the Zeiss LSM880 confocal microscope with a 63x Plan-Apochromat NA 1.4 DIC M27 oil immersion objective. Images were acquired continuously every second. Sections of supracellular actin cables were realized either parallel or perpendicular to the ablated actin cable. Perpendicular sections of cortical actin were also performed as controls.

All time-lapse experiments were conducted using the incubation chamber installed on the confocal microscopes in order to maintain a humidified atmosphere with 5 % CO2 and 37 °C throughout image acquisition. All time-lapse images from the Zeiss LSM700 and the LSM880 microscopes were acquired using Zen software.

### Western Blot

Cells were washed twice with 1X PBS and lysed in a buffer composed of 1M Tris-HCl pH 7,4 1 % SDS supplemented with 1 mM PMSF, 1 mM sodium orthovanadate, 10 µg/mL protease inhibitor cocktail (Sigma-Aldrich, #11697498001) and 50 µg/mL DNase on ice. After 30 min of incubation at 4 °C, the samples were sonicated (amplitude 40 %, 10 sec pulse on, 45 sec pulse off, 3 cycles). Lysates were then centrifuged at 1400 rpm for 30 min at 4 °C. Supernatants were collected and stored at -80 °C before use.

Protein concentrations were determined using the Pierce BCA Protein Assay Kit (Thermo Fisher Scientific, #23225). 40 µg of proteins were separated using 6 % SDS–PAGE gels and transferred to nitrocellulose membranes. Membranes were then blocked overnight in a blocking buffer composed of 1X TBS, 0,1 % Tween-20 and 5 % non-fat dry milk. The following day, they were incubated with primary antibodies diluted in the blocking buffer for 1 h at RT. Secondary antibodies, also diluted in the blocking buffer, were then applied for 1 h at RT and protein bands were revealed using a ChemiDoc imaging system from Bio-Rad.

The following primary antibodies were used: anti-p120-catenin rabbit polyclonal antibody (1:1000, Proteintech, #12180-1-AP), anti-ZO-1 mouse monoclonal antibody (1:500, Invitrogen, #740002M), anti-α-tubulin mouse monoclonal antibody (1:2000, Cell Signalling, #3873) and anti-Hsp70 mouse monoclonal antibody (1:1000, Santa Cruz Biotechnology #sc-24). The secondary antibodies were Peroxidase AffiniPure goat anti-rabbit IgG and goat anti-mouse IgG (1:5000, Jackson Immunoresearch, #111-035-144 and #115-035-062, respectively).

### Quantifications and analysis

All the following quantifications were performed using FIJI software, except the ones for laser ablation that were done using Matlab.

#### Wound area

Brightfield images showing the space between the cell sheets were used to quantify the wound area. On each image, the cell-free regions were delineated using the freehand selection tool to measure the wound area.

#### Number, elongation and area of fingers

These parameters were assessed using brightfield images of the outer periphery of the cell sheets. For each finger, a line was drawn at its base. The area of a finger was defined as the space of cells above this line. Elongation was quantified by calculating the ratio length-to-width, where width is measured across the base and length is measured from the middle of the base to the tip of the finger. To determine the number of fingers, a straight line was drawn parallel to the front of migration, spanning from one side of the image to the other. The number of fingers per mm² was calculated by dividing the number of fingers on an image by the length of this line. Only fingers with an area greater than 10 000 µm² and an elongation ratio superior to 0,33 were included in the analysis.

#### Vimentin, fibronectin and p120-catenin fluorescence intensities

To assess the mean vimentin fluorescence intensity, images of cells stained for actin and vimentin were analysed. Sum intensity projections were generated from a series of Z-planes spanning the basal to apical sides of the cells. For vimentin and fibronectin fluorescence intensity quantifications, two regions of interest (ROIs) were outlined on the actin channel using the freehand selection tool: one at the front and one at the rear of migration as shown in the figures. On the vimentin channel, the mean gray values within these ROIs were quantified. To calculate the corrected mean vimentin fluorescence intensity, the mean gray value in the background (region without cells) was substracted from the ROI measurements. The difference in vimentin fluorescence intensity between the front and the rear of migration was determined by subtracting the mean intensity at the rear from that at the front. The same procedure was applied to fibronectin measurements, substituting vimentin with fibronectin in the analysis. It was also applied to calculate the mean p120-catenin fluorescence intensity except that only one ROI outlining the entire cell area was drawn.

#### Number and area of hubs and total length of the actin cables

Cells stained for actin and an adherens junction protein (p120-catenin or β-catenin) were imaged. On each plan of the Z-stacks, the centers of hubs as well as supracellular actin cables were manually delineated using the freehand selection tool. To be considered as a hub, actin structures must meet the following criteria: they should not be on the same plane as stress fibers, should not colocalize with cell-cell junctions signals, should have a round shape with at least four ramifications and should have an area greater than 10 µm². The area of each hub was measured and the total number of hubs per mm² was calculated by quantifying the entire cell area with the freehand selection tool. To be classified as a supracellular actin cable, actin structures must fulfill these criteria: they should not be on the same plane as stress fibers, should not colocalize with cell-cell junctions signal, should be distinct from transverse actin arcs and should have a width superior to 2 µm. The total length of actin cables per mm² was determined after outlining the entire cell area.

#### Normalized fluorescence intensities in plot profiles

A line was drawn on fluorescence images along a supracellular actin cable. The plot profile tool was used to measure, on each channel, the fluorescence intensity of each pixel along this line. These values were normalized by scaling the maximum intensity in each channel to 1.

#### Kymographs

For kymograph generation, lines were drawn parallel to the direction of migration in the middle of a side cell and a leader cell using differential interference contrast (DIC) time-lapse imaging. The dynamic reslice tool was then employed to produce kymographs from these lines.

#### Cell velocities

Cell tracking of leader and side cells was performed using the manual tracking tool on DIC time-lapse imaging. Tracking was stopped once cells entered division. Cell velocities were measured using the chemotaxis and migration tool from IBIDI. The average velocity of a 4 h tracking period was plotted.

#### Actin network orientation and classification of actin structures in different orientations

Cells stained for actin were imaged and maximum intensity projections were generated from a series of apical Z-planes. If needed, images would be rotated so that their right side would coincide with the direction of migration. A mask was applied to retain pixels with values between 1500 and 15000. To quantify the orientation of actin structures, the OrientationJ Vector field plugin with the following parameters were used: a local window size of 4 pixels, a Gaussian gradient and a grid size of 10 pixels. This generates vectors with different angles and energy values depending on the orientation and signal intensity of actin structures. Only vectors with an energy value greater to 0,1 were included in the analysis and vector angles were corrected to fit within the range of -90° to 90°. The number of vectors for each angle was quantified for each image. Data from all images were combined to generate a radar chart using the Microsoft Excel software. This radar chart represents the mean orientation of actin structure with 0° representing the direction of migration. Additionally, vector angles were used to quantify, for each image, the percentage of actin structures within three different orientation categories.

#### Laser ablation

Prior to analysis, movies were processed on ImageJ and rotated so that the severed actin cable aligned with the y-axis. PIV analyses were performed using the PIVlab plugin in Matlab. If needed, masks were manually drawn on regions lacking fluorescent signal. PIV was realized using a multipass fast Fourier transform (FFT) approach with 3 passes, using an interrogation window of 64 pixels and a step size of 32 pixels. A velocity-based validation was applied on all frames.

Color-coded maps were generated for the magnitude, the vertical v-component and the horizontal u-component of junctional displacement. Areas along the actin structures that exhibited movement immediately after ablation were delineated with a rectangular region. For each parameter, data were extracted from these regions in the frames before and after ablation.

To estimate the tension release distance following ablation, two vertical lines were drawn from the ablation site to the top and bottom edges of the post-ablation frame. Along each line, magnitude values were extracted, before and after ablation. The number of cells intersected by each line was counted and divided by the total line length to estimate the average cell size, which was determined to be approximately 20 µm. To calculate the average distance to which tension was released after the ablation, data from all analyzed lines were compiled and processed using algorithms we generated on RStudio, with the assistance of ChatGPT. All the codes are available at the following address: https://github.com/ClaireBaudouin/An-extended-and-apical-supracellular-actin-network-interconnects-squamous-carcinoma-cells

### Statistical analysis

All graphical representations and statistical analyses were performed using GraphPad Prism 8.0.1 software except the charts from figure 5E drawn using Microsoft Excel software and the ones from figure 9 and Supplementary figure 5 generated using RStudio software. Each experiment was conducted independently at least three times, with the exceptions of those presented in Figure 3B and Supplementary Figure 2D. The number of independent experiments and the quantity of images or cells analysed are specified in figure legends. Given that normal distribution of the data was not formally tested, non-parametric statistical tests were employed for all analyses. For comparisons involving a single variable, we used the Mann-Whitney test for two unpaired samples, the Wilcoxon test for two paired samples and the Kruskal-Wallis test for three or more unpaired samples. For analyses involving two variables, a two-way analysis of variance (ANOVA) was applied. Information about statistical tests used for each analysis is provided in the corresponding figure legends. Values are expressed as mean ± standard deviation (s.d.) with all individual data points represented on the graphics. Statistical significance is denoted in figure legends as follows: *p < 0.05, **p < 0.01, ***p < 0.001, ****p < 0.0001. p > 0.05 was considered not significant (ns).

## Declaration of generative AI and AI-assisted technologies in the writing process

During the preparation of this work the authors used ChatGPT correct grammar and spelling. After using this tool, the authors reviewed and edited the content as needed and take full responsibility for the content of the published article.

## ABBREVIATIONS

ASAN: Apical supracellular actin network
BSA: bovine serum albumin
DIC: Differential interference contrast
DMEM: Dulbecco’s Modified Eagle Medium
EMT: Epithelial-to-mesenchymal transition
FBS: fetal bovine serum
FFT: Fast Fourier Transform
IgG: Immunoglobulin G
MAP4K4: Mitogen-activated kinase kinase kinase kinase 4
MDCK: Madin-Darby canine kidney
NMII: Non-muscle myosin II
ON: Overnight
PBS: phosphate buffered saline
PIV: Particle Image Velocimetry
pMLC2: Phosphorylated form of non-muscle myosin light chain 2
ROCK: Rho-associated kinase
ROI: Region of interest
RT: room temperature
shRNA: short-hairpin
RNA ZO-1: Zonula Occludens 1

## ACKNOWLEDGMENTS

We thank Dr Arnold Hayer (McGill University, Montréal), Dr Sébastien Carréno (IRIC, University of Montréal) and Dr Marc Therrien (IRIC, University of Montréal) for their generosity in sharing cell lines and plasmids. We also thank Christian Charbonneau from the microscopy platform (IRIC, University of Montréal), Jean-René Sylvestre and Sarah Keil for technical assistance. We thank Samuel Baudouin for his help in generating the codes for the laser ablation data analysis Finally, we thank Dr Arnold Hayer, Qiyao Lin, Dr Allen Ehrlicher and Alice Delarue (McGill University, Montréal) for the helpful discussions, and Dr Arnold Hayer for critical reading of the manuscript. We acknowledge the Fonds de recherche du Québec (FRQ) – secteur Santé for its support of the Institute for Research in Immunology and Cancer (IRIC), an FRQ-designated research center.

## REFERENCES

Alberici Delsin, L. E., Plutoni, C., Clouvel, A., Keil, S., Marpeaux, L., Elouassouli, L., Khavari, A., Ehrlicher, A. J. & Emery, G. 2023. MAP4K4 regulates forces at cell-cell and cell-matrix adhesions to promote collective cell migration. Life Sci Alliance, 6.

Amano, M., Ito, M., Kimura, K., Fukata, Y., Chihara, K., Nakano, T., Matsuura, Y. & Kaibuchi, K. 1996. Phosphorylation and activation of myosin by Rho-associated kinase (Rho-kinase). J Biol Chem, 271, 20246–9.

Arboleda-Estudillo, Y., Krieg, M., Stuhmer, J., Licata, N. A., Muller, D. J. & Heisenberg, C. P. 2010. Movement directionality in collective migration of germ layer progenitors. Curr Biol, 20, 161–9.

Balcioglu, H. E., Balasubramaniam, L., Stirbat, T. V., Doss, B. L., Fardin, M.-A., Mège, R.-M. & Ladoux, B. 2020. A subtle relationship between substrate stiffness and collective migration of cell clusters. Soft Matter, 16, 1825–39.

Bement, W. M., Forscher, P. & Mooseker, M. S. 1993. A novel cytoskeletal structure involved in purse string wound closure and cell polarity maintenance. J Cell Biol, 121, 565–78.

Bleaken, B. M., Menko, A. S. & Walker, J. L. 2016. Cells activated for wound repair have the potential to direct collective invasion of an epithelium. Mol Biol Cell, 27, 451–65.

Borghi, N., Sorokina, M., Shcherbakova, O. G., Weis, W. I., Pruitt, B. L., Nelson, W. J. & Dunn, A. R. 2012. E-cadherin is under constitutive actomyosin-generated tension that is increased at cell-cell contacts upon externally applied stretch. Proc Natl Acad Sci U S A, 109, 12568–73.

Boyer, B., Tucker, G. C., Valles, A. M., Gavrilovic, J. & Thiery, J. P. 1989. Reversible transition towards a fibroblastic phenotype in a rat carcinoma cell line. Int J Cancer Suppl, 4, 69–75.

Buckley, C. D., Tan, J., Anderson, K. L., Hanein, D., Volkmann, N., Weis, W. I., Nelson, W. J. & Dunn, A. R. 2014. Cell adhesion. The minimal cadherin-catenin complex binds to actin filaments under force. Science, 346, 1254211.

Buckley, C. E. & ST Johnston, D. 2022. Apical-basal polarity and the control of epithelial form and function. Nat Rev Mol Cell Biol, 23, 559–77.

Cai, D., Chen, S. C., Prasad, M., He, L., Wang, X., Choesmel-Cadamuro, V., Sawyer, J. K., Danuser, G. & Montell, D. J. 2014. Mechanical feedback through E-cadherin promotes direction sensing during collective cell migration. Cell, 157, 1146–59.

Chen, T., Saw, T. B., Mege, R. M. & Ladoux, B. 2018. Mechanical forces in cell monolayers. J Cell Sci, 131.

Cheng, J. C., Miller, A. L. & Webb, S. E. 2004. Organization and function of microfilaments during late epiboly in zebrafish embryos. Dev Dyn, 231, 313–23.

Chu, Y. S., Thomas, W. A., Eder, O., Pincet, F., Perez, E., Thiery, J. P. & Dufour, S. 2004. Force measurements in E-cadherin-mediated cell doublets reveal rapid adhesion strengthened by actin cytoskeleton remodeling through Rac and Cdc42. J Cell Biol, 167, 1183–94.

Clapham, D. E. 2007. Calcium signaling. Cell, 131, 1047–58.

Czerniak, N. D., Dierkes, K., D’angelo, A., Colombelli, J. & Solon, J. 2016. Patterned Contractile Forces Promote Epidermal Spreading and Regulate Segment Positioning during Drosophila Head Involution. Curr Biol, 26, 1895–901.

Danjo, Y. & Gipson, I. K. 1998. Actin ‘purse string’ filaments are anchored by E-cadherin-mediated adherens junctions at the leading edge of the epithelial wound, providing coordinated cell movement. J Cell Sci, 111, 3323–32.

Davis, M. A., Ireton, R. C. & Reynolds, A. B. 2003. A core function for p120-catenin in cadherin turnover. J Cell Biol, 163, 525–34.

De Ceccatty, M. P. 1986. Cytoskeletal organization and tissue patterns of epithelia in the sponge Ephydatia mülleri. J Morphol, 189, 45–65.

Dissanayaka, W. L., Pitiyage, G., Kumarasiri, P. V. R., Liyanage, R. L. P. R., Dias, K. D. & Tilakaratne, W. M. 2012. Clinical and histopathologic parameters in survival of oral squamous cell carcinoma. Oral Surg Oral Med Oral Pathol Oral Radiol, 113, 518–25.

Dulbecco, R., Allen, R., Okada, S. & Bowman, M. 1983. Functional changes of intermediate filaments in fibroblastic cells revealed by a monoclonal antibody. Proc Natl Acad Sci U S A, 80, 1915–8.

Engl, W., Arasi, B., Yap, L. L., Thiery, J. P. & Viasnoff, V. 2014. Actin dynamics modulate mechanosensitive immobilization of E-cadherin at adherens junctions. Nat Cell Biol, 16, 587–94.

Feng, Y., Ngu, H., Alford, S. K., Ward, M., Yin, F. & Longmore, G. D. 2013. alpha-actinin1 and 4 tyrosine phosphorylation is critical for stress fiber establishment, maintenance and focal adhesion maturation. Exp Cell Res, 319, 1124–35.

Fenteany, G., Janmey, P. A. & Stossel, T. P. 2000. Signaling pathways and cell mechanics involved in wound closure by epithelial cell sheets. Curr Biol, 10, 831–8.

Fernandez-Gonzalez, R. & Peifer, M. 2022. Powering morphogenesis: multiscale challenges at the interface of cell adhesion and the cytoskeleton. Mol Biol Cell, 33.

Goel, A., Chhabra, R., Ahmad, S., Prasad, A. K., Parmar, V. S., Ghosh, B. & Saini, N. 2012. DAMTC regulates cytoskeletal reorganization and cell motility in human lung adenocarcinoma cell line: an integrated proteomics and transcriptomics approach. Cell Death Dis, 3, e402.

Haga, H., Irahara, C., Kobayashi, R., Nakagaki, T. & Kawabata, K. 2005. Collective movement of epithelial cells on a collagen gel substrate. Biophys J, 88, 2250–6.

Hidalgo-Carcedo, C., Hooper, S., Chaudhry, S. I., Williamson, P., Harrington, K., Leitinger, B. & Sahai, E. 2011. Collective cell migration requires suppression of actomyosin at cell-cell contacts mediated by DDR1 and the cell polarity regulators Par3 and Par6. Nat Cell Biol, 13, 49–58.

Ishiyama, N., Lee, S. H., Liu, S., Li, G. Y., Smith, M. J., Reichardt, L. F. & Ikura, M. 2010. Dynamic and static interactions between p120 catenin and E-cadherin regulate the stability of cell-cell adhesion. Cell, 141, 117–28.

Ishizaki, T., Uehata, M., Tamechika, I., Keel, J., Nonomura, K., Maekawa, M. & Narumiya, S. 2000. Pharmacological properties of Y-27632, a specific inhibitor of rho-associated kinases. Mol Pharmacol, 57, 976–83.

Isozaki, Y., Sakai, K., Kohiro, K., Kagoshima, K., Iwamura, Y., Sato, H., Rindner, D., Fujiwara, S., Yamashita, K., Mizuno, K. & Ohashi, K. 2020. The Rho-guanine nucleotide exchange factor Solo decelerates collective cell migration by modulating the Rho-ROCK pathway and keratin networks. Mol Biol Cell, 31, 741–52.

Jacinto, A., Wood, W., Woolner, S., Hiley, C., Turner, L., Wilson, C., Martinez-Arias, A. & Martin, P. 2002. Dynamic analysis of actin cable function during Drosophila dorsal closure. Curr Biol, 12, 1245–50.

Kato, T., Jenkins, R. P., Derzsi, S., Tozluoglu, M., Rullan, A., Hooper, S., Chaleil, R. A. G., Joyce, H., Fu, X., Thavaraj, S., Bates, P. A. & Sahai, E. 2023. Interplay of adherens junctions and matrix proteolysis determines the invasive pattern and growth of squamous cell carcinoma. Elife, 12, e76520.

Kemp, J. P., JR. & Brieher, W. M. 2018. The actin filament bundling protein alpha-actinin-4 actually suppresses actin stress fibers by permitting actin turnover. J Biol Chem, 293, 14520–33.

Kiehart, D. P., Galbraith, C. G., Edwards, K. A., Rickoll, W. L. & Montague, R. A. 2000. Multiple forces contribute to cell sheet morphogenesis for dorsal closure in Drosophila. J Cell Biol, 149, 471–90.

Kimura, K., Ito, M., Amano, M., Chihara, K., Fukata, Y., Nakafubu, M., Yamamori, B., Feng, J., Nakano, T., Okawa, K., Iwamatsu, A. & Kaibuchi, K. 1996. Regulation of myosin phosphatase by Rho and Rho-associated kinase (Rho-kinase). Science, 273, 245–8.

Köppen, M., Fernandez, B. G., Carvalho, L., Jacinto, A. & Heisenberg, C. P. 2006. Coordinated cell-shape changes control epithelial movement in zebrafish and Drosophila. Development, 133, 2671–81.

Kovac, B., Teo, J. L., Makela, T. P. & Vallenius, T. 2013. Assembly of non-contractile dorsal stress fibers requires alpha-actinin-1 and Rac1 in migrating and spreading cells. J Cell Sci, 126, 263–73.

Laplante, C. & Nilson, L. A. 2006. Differential expression of the adhesion molecule Echinoid drives epithelial morphogenesis in Drosophila. Development, 133, 3255–64.

Lazarides, E. & Burridge, K. 1975. α-Actinin: Immunofluorescent localization of a muscle structural protein in nonmuscle cells. Cell, 6, 289–98.

Lebert, D., Squirrell, J. M., Freisinger, C., Rindy, J., Golenberg, N., Frecentese, G., Gibson, A., Eliceiri, K. W. & Huttenlocher, A. 2018. Damage-induced reactive oxygen species regulate vimentin and dynamic collagen-based projections to mediate wound repair. Elife, 7, e30703.

Lemke, S. B. & Nelson, C. M. 2021. Dynamic changes in epithelial cell packing during tissue morphogenesis. Curr Biol, 31, R1098–110.

Macpherson, I. R., Hooper, S., Serrels, A., Mcgarry, L., Ozanne, B. W., Harrington, K., Frame, M. C., Sahai, E. & Brunton, V. G. 2007. p120-catenin is required for the collective invasion of squamous cell carcinoma cells via a phosphorylation-independent mechanism. Oncogene, 26, 5214–28.

Marchiando, A. M., Graham, W. V. & Turner, J. R. 2010. Epithelial barriers in homeostasis and disease. Annu Rev Pathol, 5, 119–44.

Markov, A. G., Veshnyakova, A., Fromm, M., Amasheh, M. & Amasheh, S. 2010. Segmental expression of claudin proteins correlates with tight junction barrier properties in rat intestine. J Comp Physiol B, 180, 591–8.

Martin, A. C., Kaschube, M. & Wieschaus, E. F. 2009. Pulsed contractions of an actin-myosin network drive apical constriction. Nature, 457, 495–9.

Martin, P. & Lewis, J. 1992. Actin cables and epidermal movement in embryonic wound healing. Nature, 360, 179–83.

Maruyama, K. & Ebashi, S. 1965. Alpha-actinin, a new structural protein from striated muscle. II. Action on actin. J Biochem, 58, 13–9.

Matsui, T., Amano, M., Yamamoto, T., Chihara, K., Nakafuku, M., Ito, M., Nakano, T., Okawa, K., Iwamatsu, A. & Kaibuchi, K. 1996. Rho-associated kinase, a novel serine/threonine kinase, as a putative target for small GTP binding protein Rho. EMBO J, 15, 2208–16.

Menko, A. S., Bleaken, B. M., Libowitz, A. A., Zhang, L., Stepp, M. A. & Walker, J. L. 2014. A central role for vimentin in regulating repair function during healing of the lens epithelium. Mol Biol Cell, 25, 776–90.

Monier, B., Pelissier-Monier, A., Brand, A. H. & Sanson, B. 2010. An actomyosin-based barrier inhibits cell mixing at compartmental boundaries in Drosophila embryos. Nat Cell Biol, 12, 60–9.

Murrell, M., Oakes, P. W., Lenz, M. & Gardel, M. L. 2015. Forcing cells into shape: the mechanics of actomyosin contractility. Nat Rev Mol Cell Biol, 16, 486–98.

Omelchenko, T., Vasiliev, J. M., Gelfand, I. M., Feder, H. H. & Bonder, E. M. 2003. Rho-dependent formation of epithelial “leader” cells during wound healing. Proc Natl Acad Sci U S A, 100, 10788–93.

Ozawa, M., Baribault, H. & Kemler, R. 1989. The cytoplasmic domain of the cell adhesion molecule uvomorulin associates with three independent proteins structurally related in different species. EMBO J, 8, 1711–7.

Panfilio, K. A. & Roth, S. 2010. Epithelial reorganization events during late extraembryonic development in a hemimetabolous insect. Dev Biol, 340, 100–15.

Pedrazza, L., Schneider, T., Bartrons, R., Ventura, F. & Rosa, J. L. 2020. The ubiquitin ligase HERC1 regulates cell migration via RAF-dependent regulation of MKK3/p38 signaling. Sci Rep, 10, 824.

Peterson, L. J., Rajfur, Z., Maddox, A. S., Freel, C. D., Chen, Y., Edlund, M., Otey, C. & Burridge, K. 2004. Simultaneous stretching and contraction of stress fibers in vivo. Mol Biol Cell, 15, 3497–508.

Petitjean, L., Reffay, M., Grasland-Mongrain, E., Poujade, M., Ladoux, B., Buguin, A. & Silberzan, P. 2010. Velocity fields in a collectively migrating epithelium. Biophys J, 98, 1790–800.

Piepenhagen, P. A. & Nelson, W. J. 1993. Defining E-cadherin-associated protein complexes in epithelial cells: plakoglobin, beta- and gamma-catenin are distinct components. J Cell Sci, 104, 751–62.

Plutoni, C., Keil, S., Zeledon, C., Delsin, L. E. A., Decelle, B., Roux, P. P., Carreno, S. & Emery, G. 2019. Misshapen coordinates protrusion restriction and actomyosin contractility during collective cell migration. Nat Commun, 10, 3940.

Poujade, M., Grasland-Mongrain, E., Hertzog, A., Jouanneau, J., Chavrier, P., Ladoux, B., Buguin, A. & Silberzan, P. 2007. Collective migration of an epithelial monolayer in response to a model wound. Proc Natl Acad Sci U S A, 104, 15988–93.

Ramel, D., Wang, X., Laflamme, C., Montell, D. J. & Emery, G. 2013. Rab11 regulates cell-cell communication during collective cell movements. Nat Cell Biol, 15, 317–24.

Reffay, M., Parrini, M. C., Cochet-Escartin, O., Ladoux, B., Buguin, A., Coscoy, S., Amblard, F., Camonis, J. & Silberzan, P. 2014. Interplay of RhoA and mechanical forces in collective cell migration driven by leader cells. Nat Cell Biol, 16, 217–23.

Reffay, M., Petitjean, L., Coscoy, S., Grasland-Mongrain, E., Amblard, F., Buguin, A. & Silberzan, P. 2011. Orientation and polarity in collectively migrating cell structures: statics and dynamics. Biophys J, 100, 2566–75.

Reynolds, A. B., Daniel, J., Mccrea, P. D., Wheelock, M. J., Wu, J. & Zhang, Z. 1994. Identification of a new catenin: the tyrosine kinase substrate p120cas associates with E-cadherin complexes. Mol Cell Biol, 14, 8333–42.

Rimm, D. L., Koslov, E. R., Kebriaei, P., Cianci, C. D. & Morrow, J. S. 1995. Alpha 1(E)-catenin is an actin-binding and -bundling protein mediating the attachment of F-actin to the membrane adhesion complex. Proc Natl Acad Sci U S A, 92, 8813–7.

Rodriguez-Diaz, A., Toyama, Y., Abravanel, D. L., Wiemann, J. M., Wells, A. R., Tulu, U. S., Edwards, G. S. & Kiehart, D. P. 2008. Actomyosin purse strings: renewable resources that make morphogenesis robust and resilient. HFSP J, 2, 220–37.

Röper, K. 2012. Anisotropy of Crumbs and aPKC drives myosin cable assembly during tube formation. Dev Cell, 23, 939–53.

Summerbell, E. R., Mouw, J. K., Bell, J. S. K., Knippler, C. M., Pedro, B., Arnst, J. L., Khatib, T. O., Commander, R., Barwick, B. G., Konen, J., Dwivedi, B., Seby, S., Kowalski, J., Vertino, P. M. & Marcus, A. I. 2020. Epigenetically heterogeneous tumor cells direct collective invasion through filopodia-driven fibronectin micropatterning. Sci Adv, 6.

Tamada, M., Perez, T. D., Nelson, W. J. & Sheetz, M. P. 2007. Two distinct modes of myosin assembly and dynamics during epithelial wound closure. J Cell Biol, 176, 27–33.

Thomas, W. A., Boscher, C., Chu, Y. S., Cuvelier, D., Martinez-Rico, C., Seddiki, R., Heysch, J., Ladoux, B., Thiery, J. P., Mege, R. M. & Dufour, S. 2013. alpha-Catenin and vinculin cooperate to promote high E-cadherin-based adhesion strength. J Biol Chem, 288, 4957–69.

Tunggal, J. A., Helfrich, I., Schmitz, A., Schwarz, H., Gunzel, D., Fromm, M., Kemler, R., Krieg, T. & Niessen, C. M. 2005. E-cadherin is essential for in vivo epidermal barrier function by regulating tight junctions. EMBO J, 24, 1146–56.

Vaezi, A., Bauer, C., Vasioukhin, V. & Fuchs, E. 2002. Actin Cable Dynamics and Rho/Rock Orchestrate a Polarized Cytoskeletal Architecture in the Early Steps of Assembling a Stratified Epithelium. Developmental Cell, 3, 367–381.

Van Straaten, H. W. M., Sieben, I. & Hekking, J. W. M. 2002. Multistep role for actin in initial closure of the mesencephalic neural groove in the chick embryo. Dev Dyn, 224, 103–8.

Vicente-Manzanares, M., Ma, X., Adelstein, R. S. & Horwitz, A. R. 2009. Non-muscle myosin II takes centre stage in cell adhesion and migration. Nat Rev Mol Cell Biol, 10, 778–90.

Volberg, T., Geiger, B., Kartenbeck, J. & Franke, W. W. 1986. Changes in membrane-microfilament interaction in intercellular adherens junctions upon removal of extracellular Ca2+ ions. J Cell Biol, 102, 1832–42.

Walker, J. L., Wolff, I. M., Zhang, L. & Menko, A. S. 2007. Activation of SRC kinases signals induction of posterior capsule opacification. Invest Ophthalmol Vis Sci, 48, 2214–23.

Walker, J. L., Zhai, N., Zhang, L., Bleaken, B. M., Wolff, I., Gerhart, J., George-Weinstein, M. & Menko, A. S. 2010. Unique precursors for the mesenchymal cells involved in injury response and fibrosis. Proc Natl Acad Sci U S A, 107, 13730–5.

Wang, H., Guo, X., Wang, X., Wang, X. & Chen, J. 2020. Supracellular Actomyosin Mediates Cell-Cell Communication and Shapes Collective Migratory Morphology. iScience, 23, 101204.

Williams-Masson, E. M., Malik, A. N. & Hardin, J. 1997. An actin-mediated two-step mechanism is required for ventral enclosure of the C. elegans hypodermis. Development, 124, 2889–901.

Wood, W., Jacinto, A., Grose, R., Woolner, S., Gale, J., Wilson, C. & Martin, P. 2002. Wound healing recapitulates morphogenesis in Drosophila embryos. Nat Cell Biol, 4, 907–12.

Yamaguchi, N., Mizutani, T., Kawabata, K. & Haga, H. 2015. Leader cells regulate collective cell migration via Rac activation in the downstream signaling of integrin beta1 and PI3K. Sci Rep, 5, 7656.

Yao, M., Qiu, W., Liu, R., Efremov, A. K., Cong, P., Seddiki, R., Payre, M., Lim, C. T., Ladoux, B., Mege, R. M. & Yan, J. 2014. Force-dependent conformational switch of alpha-catenin controls vinculin binding. Nat Commun, 5, 4525.

Yonemura, S. 2017. Actin filament association at adherens junctions. J Med Invest, 64, 14–19.

Yonemura, S., Wada, Y., Watanabe, T., Nagafuchi, A. & Shibata, M. 2010. alpha-Catenin as a tension transducer that induces adherens junction development. Nat Cell Biol, 12, 533–42.

Zihni, C., Mills, C., Matter, K. & Balda, M. S. 2016. Tight junctions: from simple barriers to multifunctional molecular gates. Nat Rev Mol Cell Biol, 17, 564–80.

