## Supplemental figures for "A supracellular actin network transmits forces over long distances at the apical surface of squamous carcinoma cells"

Supplementary figure 1

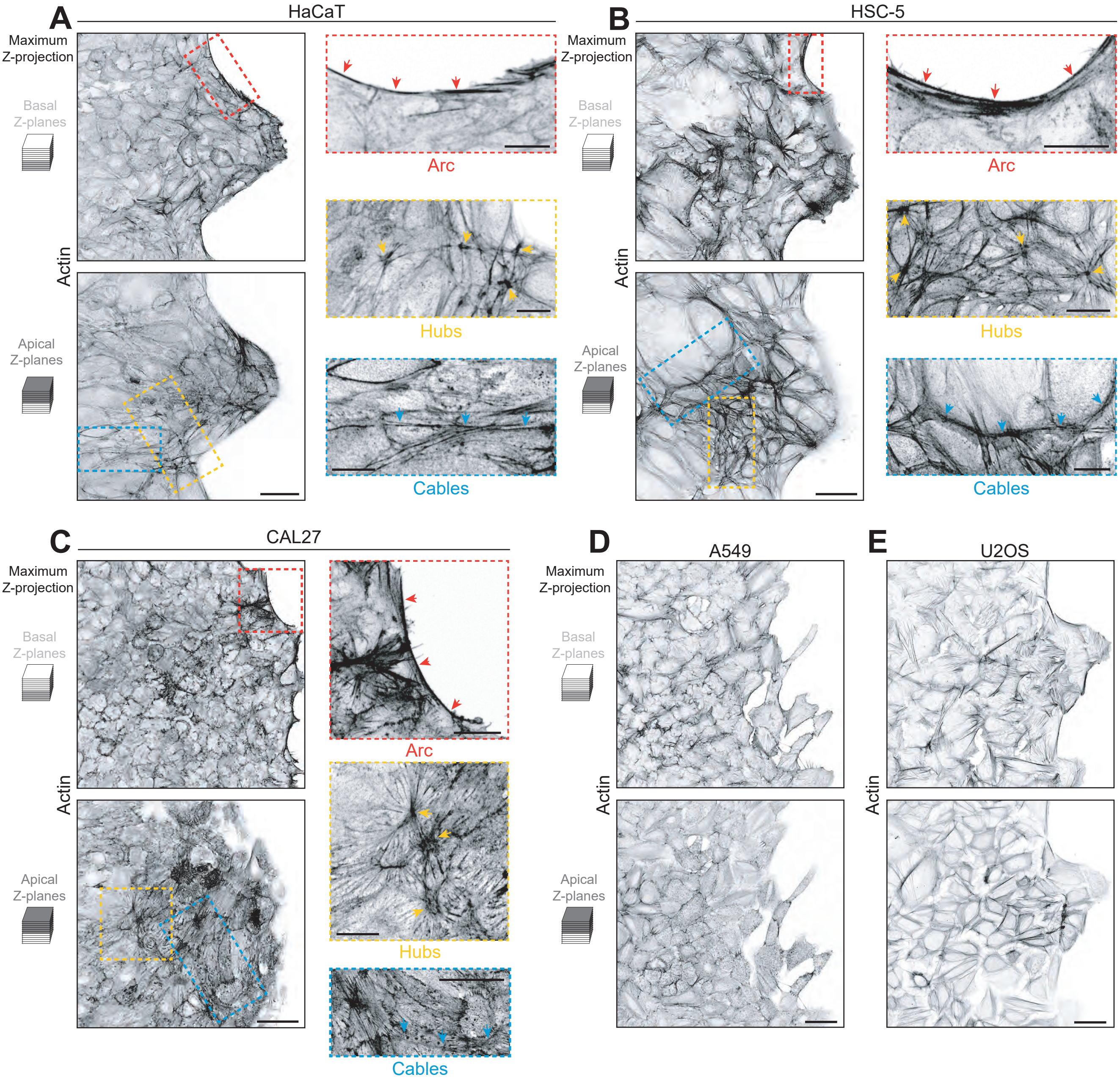

### Supplementary figure 2

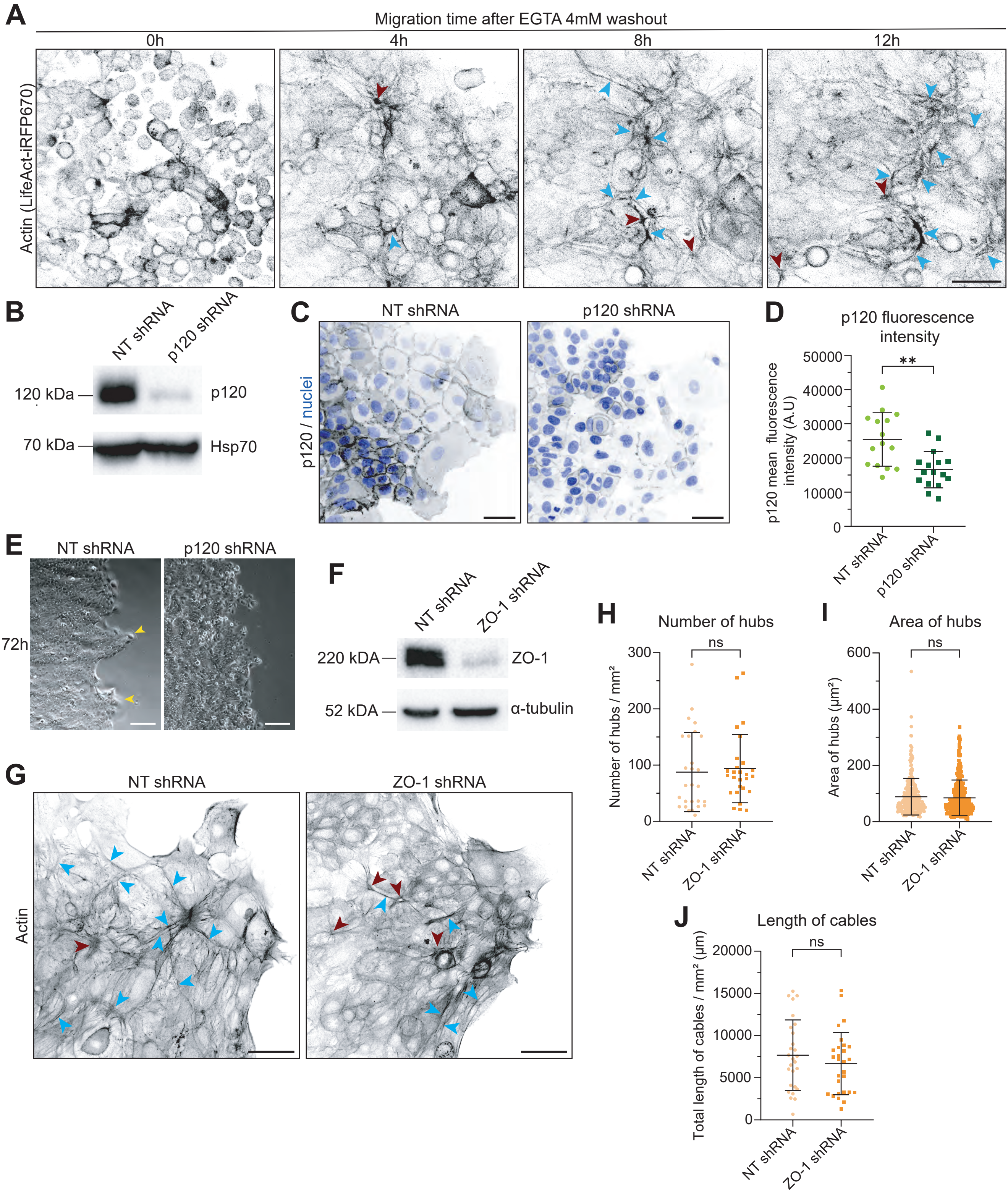

Supplementary figure 3

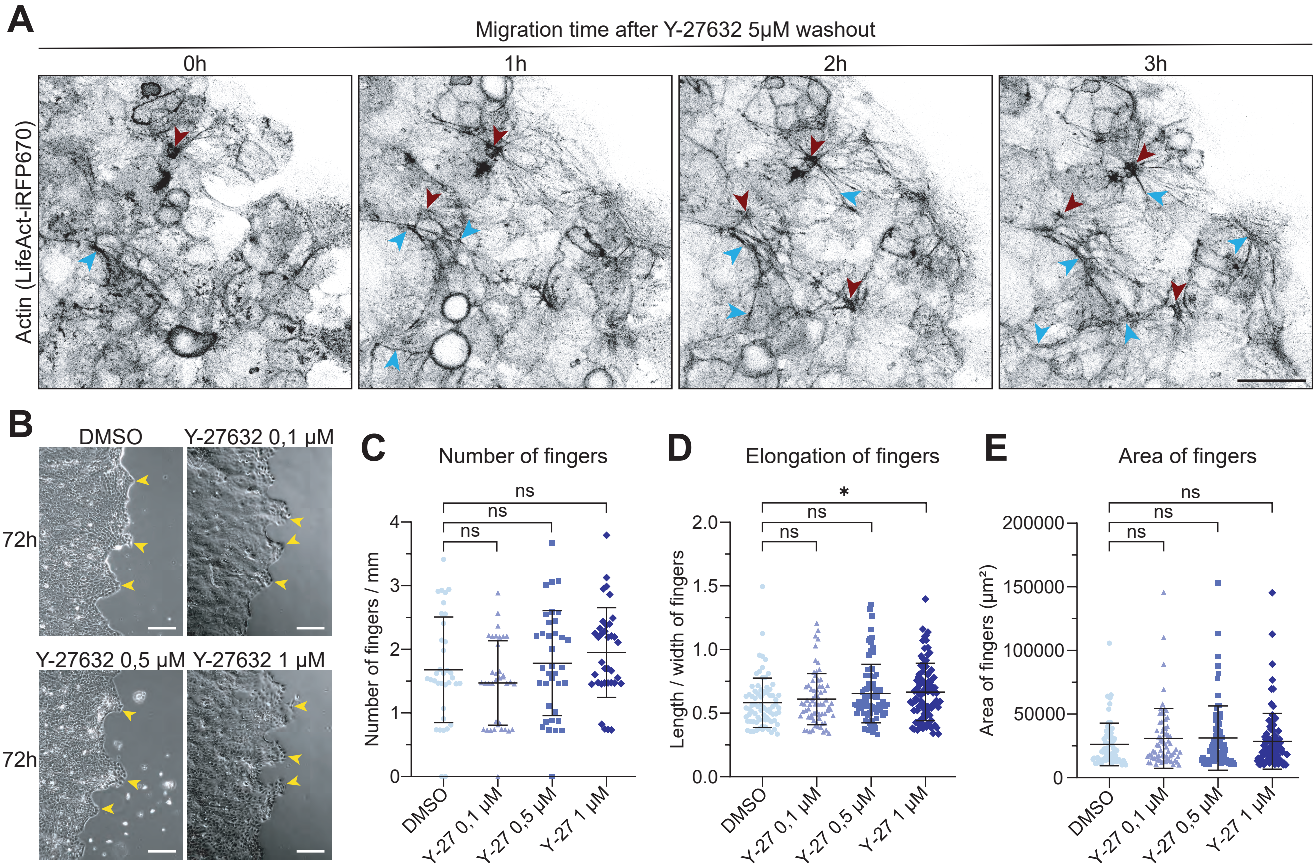

Supplementary figure 4

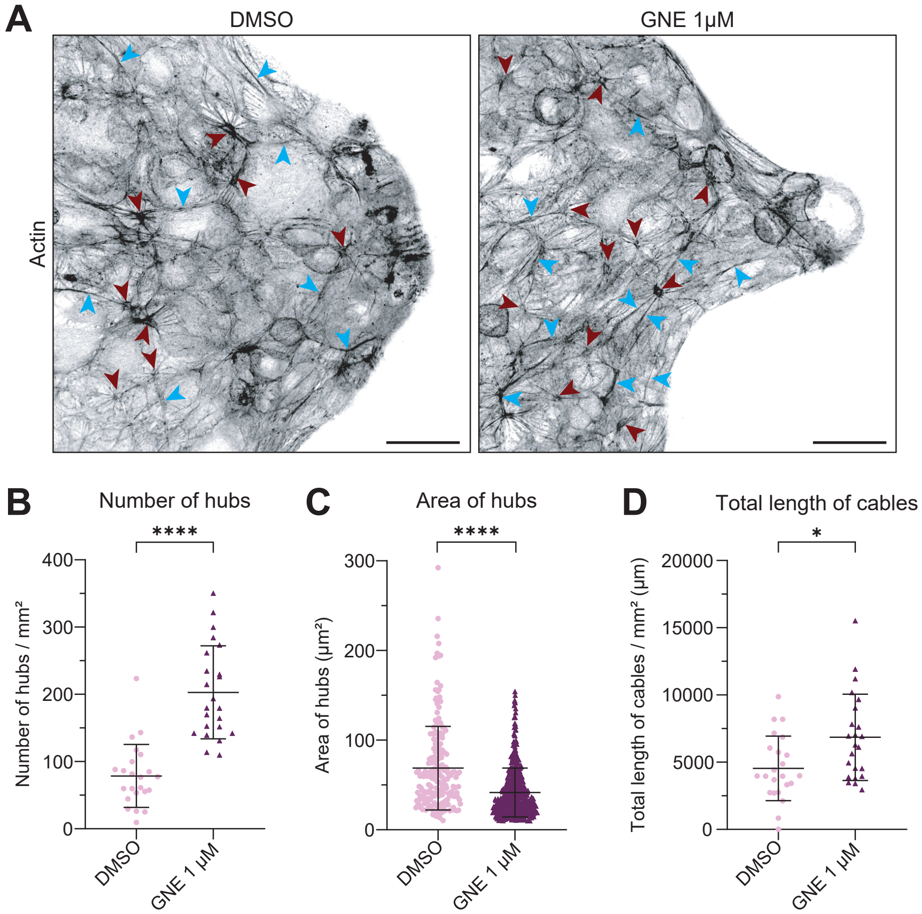

Supplementary figure 5

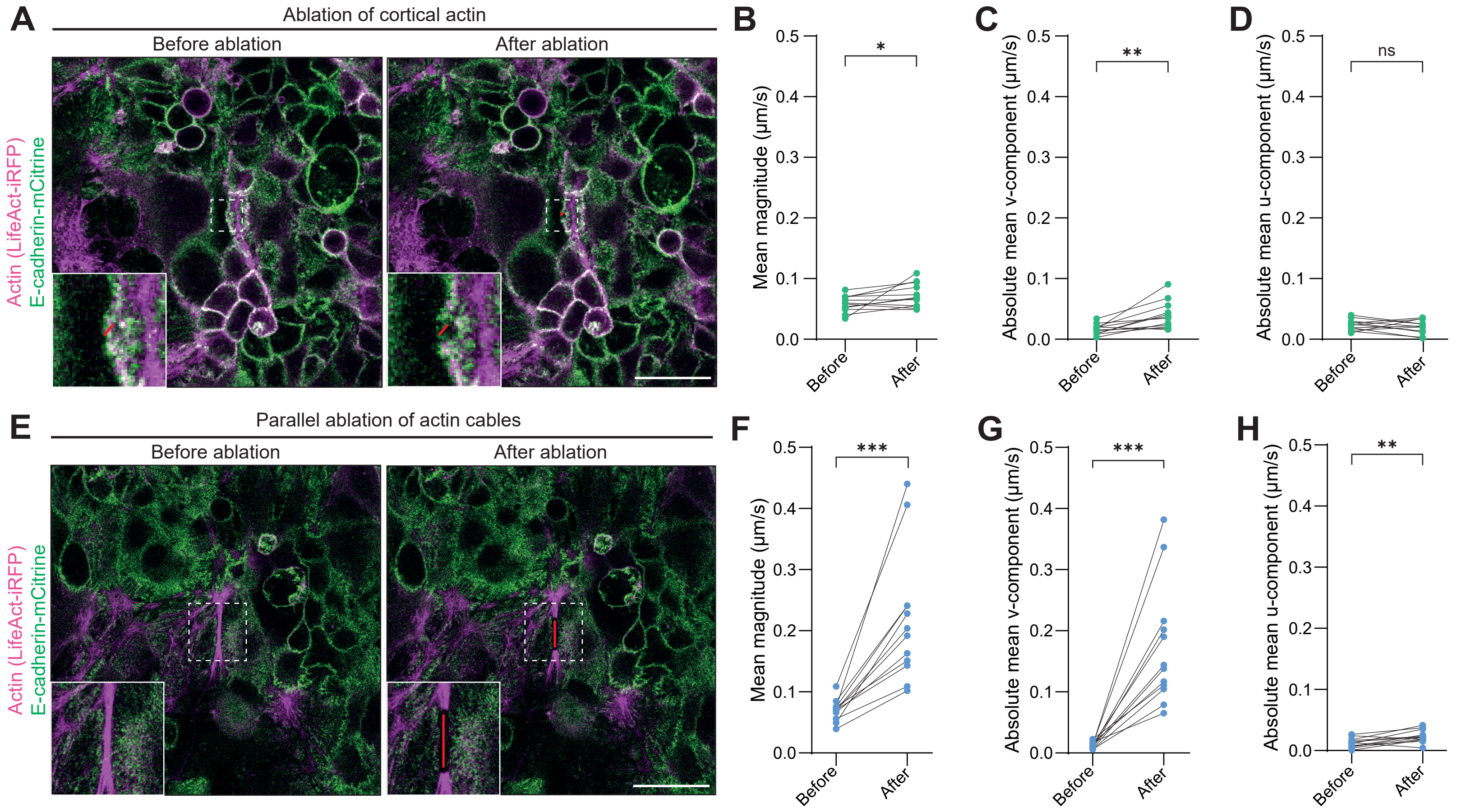
